# scLongTree: an accurate computational tool to infer the longitudinal tree for single-cell DNA sequencing data

**DOI:** 10.1101/2023.11.11.566680

**Authors:** Rituparna Khan, Pankaj Bhattarai, Liting Zhang, Xin Maizie Zhou, Xian Mallory

## Abstract

Longitudinal single-cell DNA sequencing (scDNA-seq) refers to single-cell data sequenced at different time points providing more knowledge of the order of mutations than scDNA-seq taken at only one time point. The technique can facilitate the inference of subclonal trees that depict the evolution of cancer cells and facilitate understanding of how cancer grows, with implications for prognosis and treatment. There is currently a scarcity of tools that can infer subclonal trees based on longitudinal scDNA-seq, and existing tools are limited in accuracy and scale. We therefore introduce scLongTree, a computational tool that can accurately infer a subclonal tree based on longitudinal scDNA-seq. ScLongTree is scalable to hundreds of mutations, and outperforms state-of-the-art tools such as LACE, SCITE, and SiCloneFit on a comprehensive simulated dataset. Tests on a real dataset, SA501, showed that scLongTree can more accurately interpret the progressive growth of the tumor than LACE, and is more robust to different numbers of mutations being used. Tests on a large AML dataset AML107, which has 4,617 cells, show that scLongTree is scalable to thousands of cells. ScLongTree is freely available on https://github.com/compbio-mallory/sc_longitudinal_infer.

**Key points:**

- We propose scLongTree that can infer the subclonal longitudinal tree for cancer given single-cell DNA sequencing data, and thus can facilitate the study of cancer evolution given the dataset from multiple time points.
- Multiple simulated data show that scLongTree is more accurate than existing state-of-the-art methods such as LACE, SCITE and SiCloneFit.
- ScLongTree has been shown to have a higher scalability than LACE and thus can be applicable to the datasets that have hundreds of mutations.
- The experiment on a real SA501 shows that scLongTree is more robust to the number of mutations than LACE. It consistently generates the same longitudinal tree even under different sets of mutations.
- The experiment on AML107 shows that scLongTree is scalable to thousands of cells.

## 1 Introduction

Cancer develops by acquiring somatic mutations. Different sets of cancer cells may gain different sets of mutations. This phenomenon, known as intra-tumor heterogeneity (ITH), confounds cancer treatment, prognosis, and prevention of metastasis [15, 2, 24, 16]. Unraveling the evolutionary history of tumor cells helps to characterize ITH and thus facilitates the design of the treatment plan. Such an evolutionary history can be characterized by a subclonal tree, the nodes of which represent the subclones of cells, with each subclone having a unique set of mutations. The edges of the tree represent the parent-child relationship [30] where mutations are accumulated. Thus the subclonal tree is a rooted directed tree, where the root represents the first normal cell that has not yet gained any somatic mutations.

Unlike the traditional “bulk” sequencing that combines millions of cells together and thus cannot tell which mutation belongs to which cell, single-cell DNA sequencing (scDNA-seq) makes it possible to separately sequence each cell [19, 17], providing a high resolution of subclonality and allowing a more accurate reconstruction of the subclonal tree [28]. However, scDNA-seq may be erroneous, including false positive (FP) mutations, false negative (FN) mutations and missing entries due to uneven amplifications and low coverage, in addition to doublet errors [28]. Since the advent of scDNA-seq, there have been multiple attempts to decipher ITH using scDNA-seq data which involved necessary steps to overcome these errors, such as SCITE [10], OncoNEM [22], SASC [4], Sifit [27] and SiCloneFit [29]. These state-of-the-art computational tools typically assume that the data collected were sequenced at one time point, and thus do not consider the case when the data were collected at multiple time points, even when the time points are known for each cell sequenced.

Recently, scDNA-seq datasets that were collected from different time points have emerged, including whole exome or targeted sequencing [20, 31], patient-derived tumor xenograft models (PDXs) [5, 8], as well as datasets that were collected pre-, mid- and post-treatment [12], or before and after relapse [18]. To reconstruct the evolutionary history given such longitudinal scDNA-seq data, new computational tools considering the time points when each cell was sequenced are needed. Considering time points helps to 1) resolve the mutation order, especially when there are parallel or back mutations involved; 2) infer the unobserved nodes in between two time points; 3) tracking clonal dynamics. More details of the advantages of considering time points with illustrations can be found in **Supplemental Section 1** and **Supplemental Figs. 1-3**. To our knowledge, LACE [21] is the only computational tool that is specifically designed for inferring a subclonal tree given longitudinal scDNA-seq data, in which each node represents a genotype that is contained by a set of cells, and each edge represents either a parental or a persistence relationship (i.e. parent and child having the same genotype). Such a tree was called “longitudinal subclonal tree” by LACE [21] since it incorporates the time points each cell was sequenced. Specifically, LACE models the problem of inferring the longitudinal subclonal tree by seeking the solution to a weighted maximum likelihood problem that allows distinct error rates across different time points, whereas the weight at each time point can be tuned by the prior knowledge of the quality of the data, as well as the number of cells at each time point. At each time point, LACE searches for the genotype each cell contains and the mutations each genotype has, the latter of which was called “genotype-mutation” matrix and is shared by all time points. Since LACE holds the infinite site assumption (ISA) [13, 9], there is a one to one correspondence between the “genotype-mutation” matrix and the tree, whereas the tree corresponding to the “genotype-mutation” matrix is a perfect phylogeny [7]. However, ISA disallows parallel mutations (mutations that occur in more than one edge) and back mutations (mutations that were gained and then lost) [14], the latter of which often occur due to loss of copies in the genome [27]. LACE searches for the optimal solution by Monte Carlo Markov Chain (MCMC) to solve a Boolean matrix factorization problem. However, LACE is limited in its scalability by the number of mutations, and thus cannot render any tree when the number of mutations of interest is in the hundreds. Moreover, the genotypes present in LACE’s tree have to be represented by at least one cell. In other words, LACE’s methodology cannot reconstruct a subclone that once existed in the cancer evolution history but was never sequenced. However, such reconstruction of the unobserved subclones is important because when two sequencing time points were far from each other, many events occurred between the two time points and it is possible that a subclone was not sequenced before evolving and branching into other subclones.

We developed scLongTree, a computational tool to infer the longitudinal subclonal tree based on longitudinal scDNA-seq data from multiple time points. In contrast to LACE [21], scLongTree uses a combinatorial approach to infer the longitudinal tree and a *k* -Dollo model [6] to correct for parallel and back mutations after the tree is constructed and the cells are clustered and placed on the tree nodes. In addition, scLongTree is scalable to hundreds of mutations and thousands of cells and can complete the inference within a couple of hours. Moreover, scLongTree has been designed to be able to reconstruct unobserved subclones that are not represented by any cells sequenced. Most importantly, by adopting several statistical methods as well as corroborating events across distinct time points, scLongTree is able to identify spurious subclones and eliminate them, and thus has a high accuracy in the tree reconstruction and mutation and cell placement. We conducted comprehensive simulation and real dataset experiments to test the accuracy, robustness and computational resources of scLongTree, and found that scLongTree outperformed existing state-of-the-art methods such as LACE, SCITE, SiCloneFit, RobustClone and SCG.

## 2 Methods

As the input, we assume the mutations have been detected on each cell. The detection can be done by the state-of-the-art methods such as Monovar [26] or GATK HaplotypeCaller [25]. Given a cell by mutation matrix in which each cell’s time point is known (**Fig. 1A**), scLongTree starts with clustering the cells using an existing tool BnpC [1] at each time point (**Fig. 1B**). Note that our input data is longitudinal, with each cell associated with a specific time point. This distinguishes our approach from traditional tree algorithms such as SCITE, SiFit, SiCloneFit, which do not model temporal structure and instead treat all cells as if they originate from the same time point. **Supplemental Section 1** listed more details of the difference in terms of inference and input data between the two and the advantages of considering time points while inferring the tree. ScLongTree then ranks all clusters by their probability, and from the cluster with the lowest probability to the highest, it checks whether the cluster is a spurious cluster (**Fig. 1C**). For each potentially spurious cluster, scLongTree infers a longitudinal subclonal tree and calculates its probability after removing the cluster and ascribing its cells to other clusters (**Fig. 1D**). Such an inference of the tree is based on the knowledge of the inferred underlying genotype of each cluster and the time point of each cluster. It then decides that the potentially spurious cluster is truly spurious and should be eliminated if it satisfies the following two conditions. First, the removal of the cluster will increase the probability of the tree. Second, the removal of the cluster will not lead to an excessive increase of the FP or FN rate. Once all clusters are checked and a tree is inferred, scLongTree corrects parallel and back mutations according to the *k* -Dollo model (**Fig. 1E**) and infers the final FP and FN rates. This correction does not change the tree structure retrospectively, but aims to refine mutation assignments on the inferred tree by enforcing bounded recurrence and loss events according to the *k* -Dollo assumption. Finally, since different BnpC runs may result in different initial clustering and correspondingly different longitudinal subclonal trees, we select the best longitudinal subclonal tree based on a combination of the final FP and FN rates and the probability of the tree from multiple BnpC runs. We list the algorithm of the whole pipeline in **Supplemental Algorithm S1**. In the following, we explain each step in more detail.

**Figure 1:**
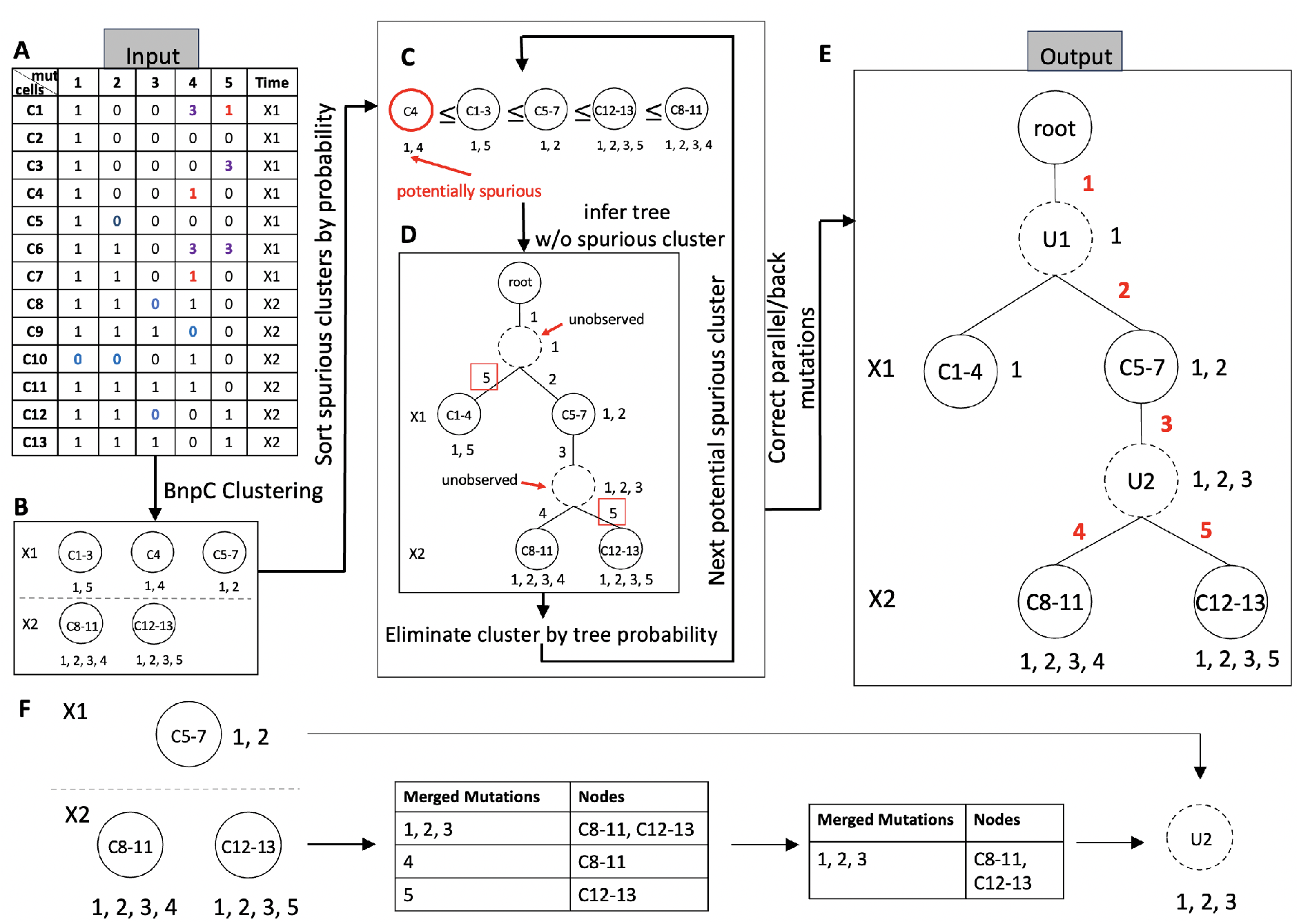
Overview of scLongTree. **A.** Input is a cell by mutation matrix, the last column of which contains the time point for each cell. Observed data may have False Positive (FP) SNVs (red “1”s), False Negative (FN) SNVs (blue “0”s) and missing entries (“3”s). **B**. ScLongTree first runs an existing cell clustering tool BnpC to cluster cells at each time point. **C**. It then sorts all subclones by their probabilities calculated using **Eq. 1**. Here the subclone with the lowest probability is the subclone with cell C4, a potentially spurious subclone. **D**. From the cluster with the lowest probability to the one with the highest probability, using the proposed combinatorial tree algorithm, scLongTree infers a longitudinal cluster tree without the potentially spurious cluster, supposing the potential spurious cluster is eliminated. The resulting tree’s probability and inferred FP/FN rates will decide whether the cluster is truly spurious and shall be eliminated. Such a process iterates between **C** and **D** until all potentially spurious clusters are checked for elimination. In the tree shown here, inside each node annotates the cells belonging to this cluster, and next to each node annotates the mutations that this cluster carries. Dashed circles represent unobserved nodes. On the edges annotates the mutations. Here mutation “5” in red boxes has been annotated on two edges, showing that it is potentially a parallel mutation. **E**. ScLongTree then corrects parallel and back mutations in concordance to the *k* -Dollo model. Mutation “5” is annotated on only one edge after this step. **F**. A detailed illustration of adding an unobserved subclone. Given the subclones in two consecutive time points *p* and *p* + 1, this algorithm searches for the largest set of mutations shared by the largest possible set of subclones in time point *p* + 1. By comparing this set of shared mutations with the mutations in time point *p*, an unobserved node can be constructed.

### 2.1 Clustering cells

ScLongTree’s first step is to cluster cells at each time point (**Fig. 1B**). We utilize an existing state-of-the-art Bayesian-based method, BnpC [1], to perform an initial cell clustering. BnpC models the observed genotypes of single cells using a probabilistic graphical model in which the false positive and false negative rates are latent variables. These rates, together with the underlying genotype of each cluster and the assignment of cells to clusters, determine the observed genotypes. BnpC estimates the latent variables using MCMC to maximize the full posterior distribution. The method assumes that cells within a subclone share the same genotype, aside from errors due to false positives and false negatives. BnpC is scalable to thousands of cells and its accuracy is overall advantageous compared with other cell clustering methods such as SCG and SCClone [11].

We run BnpC at each time point instead of clustering cells across all time points together because the former renders a few advantages. First, each time point has its own false positive and false negative rates, and BnpC can infer them independently. Second, the subclones that differ by a small number of mutations but are distributed in two different time points will not be mis-clustered together. In our simulation (**Section 3.1**), we further investigated scLongTree’s clustering accuracy by running BnpC pooling and not pooling cells across all time points, respectively. We found that the difference is negligible. Nevertheless, users can leverage prior knowledge to decide whether rare subclones are expected across time points, since pooling all cells may help capture sparse cells from rare subclones that are distributed across multiple time points.

Since BnpC is a Bayesian method and is not deterministic, we run BnpC five times and obtain five clustering results for each time point. A detailed analysis investigating BnpC’s stability is in **Section 3.1**.

### 2.2 Eliminating non-*bona fide* clusters and refining the longitudinal subclonal tree

We found that BnpC may erroneously over-cluster the cells and the clusters that are composed of a small number of cells are very likely a result of over-clustering. We aim to identify the clusters that are not *bona fide* and eliminate them (**Fig. 1C**). The whole identification and elimination are done in a probabilistic manner considering the whole tree structure.

First, based on BnpC’s inferred FP and FN rates, and the inferred underlying genotype for each cluster at each time point, we infer the posterior probability that the cluster is a *bona fide* cluster instead of a result of false clustering. A cluster’s probability shall be high if the cluster is composed of a large number of supporting cells, or if the cells inside it highly agree with each other on their genotypes and do not have many missing entries. Mathematically, suppose there are *v* cells that are in subclone *l* at time point *p*. Suppose there are in total *m* mutations involved in the analysis in all time points. Let *g* be the consensus genotype inferred from BnpC for subclone *l*, and *g* = (*g*_1_, *g*_2_, …, *g*_*m*_). *g*_*j*_ ∈ {0, 1}, *j* = 1, …, *m*, in which 0 and 1 represent absence and presence of the mutations, respectively. Let the observed data for these cells be *D*_1_, *D*_2_, …, *D*_*v*_, each of which is a vector composed of *m* mutations. Thus, *D*_*ij*_ represents the observed genotype for mutation *j* at cell *i* that was assigned to subclone *l. D*_*ij*_ ∈ {0, 1, 3}, *i* = 1, …, *v, j* = 1, …, *m*, in which 0, 1 and 3 represent absence, presence and missing of the mutations, respectively. We calculate the probability of the consensus genotype *g* given the data *D*_1_, *D*_2_, …, *D*_*v*_ from the *v* cells belonging to this subclone in **Eq. 1** by the Bayes theorem and the assumption that each mutation is independent from each other.

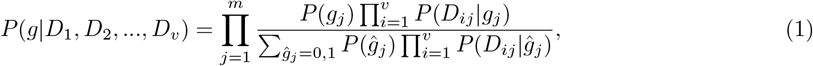

Here *P* (*D*_*ij*_|*g*_*j*_) is a likelihood that is defined in **Eq. 2**. In **Eq. 2**, *α, β* and *γ* are the initially estimated FP, FN and missing rate inferred from BnpC at time point *l*. The prior probabilities for the consensus genotype *P* (*g*_*j*_) in **Eq. 1** is defined as (1−*γ*)*/*2 for both *g*_*j*_ = 1 and *g*_*j*_ = 0. This choice reflects a minimally informative assumption: in the absence of additional information, a mutation is equally likely to be present or absent, while the remaining probability *γ* accounts for missing data and the associated uncertainty. Including *γ* ensures that the likelihood in **Eq**. 2 sums to 1 across all possible states of *g*_*j*_ and also provides a natural generalization for scenarios where the prior probabilities for *g*_*j*_ = 0 and *g*_*j*_ = 1 may differ in the future.

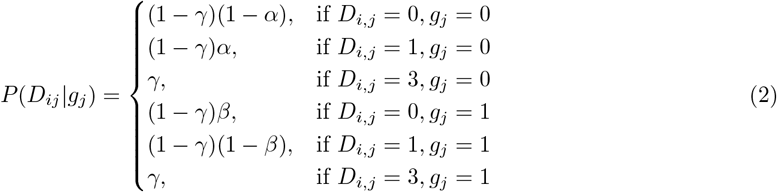

Given the posterior probability for each cluster, we increasingly sort the clusters by their posterior probability across all time points. In a serial manner, for each cluster, we examine whether it is subject to be eliminated by several criteria. The first criterion is whether the elimination of the cluster will still result in reasonable FP and FN rates such that they are smaller than a threshold. We avoid setting a hard threshold but let our algorithm learn the threshold from the data, the detail of which is described in **Supplemental Section 3**.

Notice that the elimination of a cluster may result in inflated FP or FN rate because of the reassignment of the cells that originally belong to this cluster to other clusters in the same time point (**Supplemental Section 5**). We use FP and FN rates as our quality control because high FP and/or FN values are indicators of false clustering results. Our second criterion is that the tree support score after the elimination of the cluster shall increase by any amount. We define the tree support score as the product of the posterior probabilities of all nodes in the tree. We infer the longitudinal subclonal tree before and after the elimination of the cluster by our tree algorithm described in **Sec. 2.3**. Such a frequent inference of the longitudinal subclonal tree takes advantage of our fast tree inference algorithm which typically takes several seconds for a run. If a cluster passes both criteria, we eliminate the cluster. The resulting cells will be reassigned to the other clusters in the same time point by the maximum likelihood using **Eq. 3**. More details of how we reassign the cells of the eliminated cluster to other clusters can be found in **Supplemental Section 5**.

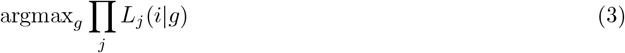

in which *L*_*j*_(*i*|*g*) is defined in **Eq. 4**.

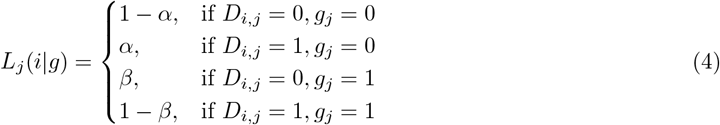

If a cluster is eliminated, we update the posterior probabilities and the order of all clusters across all time points for the next round of elimination. This process iterates until all clusters have been checked (**Fig. 1C, D**). Notice that each time we eliminate a spurious cluster, we will re-infer the tree using the algorithm described in **Sec. 2.3**. Correspondingly, the FP and FN rate will be updated based on the new tree structure when a spurious cluster is eliminated.

### 2.3 Inferring a longitudinal subclonal tree

When we build our longitudinal subclonal tree, each cluster represents a node in the tree. Since our whole pipeline requires frequent re-construction of the tree topology, we design our tree inference algorithm such that it is light-weighted. Since it is possible that two consecutive sequencing time points are far from each other and thus miss the important branching point that occur in between the two sequencing points, our tree inference algorithm has been designed to be able to detect unobserved nodes to catch the important branching point that are missed in the sequenced data. Since parallel mutations are not frequent, whereas back mutations are more likely to happen due to the copy number losses, we build our tree following the *k* -Dollo model, which does not allow parallel mutations but *k* back mutations. The following describes our tree inference algorithm in detail.

First, we link the nodes that share the same consensus genotypes between two consecutive time points as it has been observed that the subclones may persist from one time point to another. The remaining nodes may either connect to a node from the previous time point, or to an unobserved node that lies between the two consecutive time points. Here an unobserved node is defined as a node that occurs in between two neighboring time points and has different genotypes from any of the node in these two time points. It is essential to infer a tree with the unobserved nodes because the two sequencing time points might be too far away to catch all branching points, leaving some important nodes not observed. Inferring unobserved nodes is a novel feature in scLongTree, which existing methods such as LACE does not have. Our algorithm of inferring an unobserved node is not probabilistic but combinatorial to avoid large search space and long search time, facilitating frequent tree inference in the step of eliminating spurious clusters. In the following, we describe in more detail how we infer unobserved node with which we can construct the whole tree’s topology. Since it is possible that back mutations have to be introduced when adding an unobserved node, we describe our algorithm for two scenarios, which are the case when there is no back mutation introduced and the case when there are back mutations that have to be introduced.

Since an unobserved node’s genotype is not present in the clustered nodes, here we design an algorithm to identify the unobserved nodes, link them to the observed nodes, and infer their genotypes (**Fig. 1F**). Such an algorithm shall consider the genotypes of the nodes for the two time points *p* and *p* + 1, and insert the unobserved nodes in between these two time points so that the parallel mutations and back mutations placed in between time points *p* and *p* + 1 are as few as possible. Our algorithm will have multiple rounds of iterations, each round aiming to find the largest set of mutations contained by the largest possible set of nodes in time point *p* + 1. Here we describe our algorithm to come up with only one layer of unobserved nodes between time points *p* and *p* + 1. In **Supplemental Section 4**, we give a more detailed algorithm that can insert multiple layers of unobserved nodes between time points *p* and *p* + 1.

Suppose every cluster in the tree is called a node. At time point *p* + 1 (X2 in **Fig. 1F**), we build a hash table whose key is a mutation that appear in any of the nodes in time *p* + 1 and value is a set of the nodes at time *p* + 1 that contain this mutation. We then merge the keys that have the same value. For example, in **Fig. 1F**, mutations 1, 2, and 3 are merged because they are contained by the same set of nodes, C8-11, and C12-13. This step helps us to quickly find out which set of mutations is shared by the largest number of clusters. The unobserved nodes in between time points *p* and *p* + 1 are supposed to connect two or more than two nodes in time *p* + 1, forming a tree structure. Thus if a key has only one node in the value, the key will be eliminated for forming an unobserved node. Thus mutations 4 and 5 in **Fig. 1F** are removed because they appear only in one node. If a key is exactly the same as the consensus genotype of a node in time point *p*, then there is no need to add such an unobserved node. We eliminate such keys as well. Next, we sort the remaining key-value pairs in a reverse order. In **Fig. 1F**, there is only one pair left, which is ({1, 2, 3}, {C8-11, C12-13}). We then check for each pair, whether a key is a proper superset of the consensus genotype of a node in time point *p*. This is the step to decide whether back mutations will be introduced between the nodes in time point *p* and the unobserved nodes. If the set of mutations inside the key is a superset of the set of the mutations of a node, say node *A*, in time point *p*, we add an unobserved node in between time points *p* and *p* + 1, whose mutations are those in the key. We add an edge connecting the unobserved node with node *A* in time point *p*. We also add edges connecting the unobserved with the nodes in the values corresponding to the key. Here in **Fig. 1F**, an unobserved node U2 is added whose mutation set is {1, 2, 3}. U2 will then be connected with node C5-7, gaining a new mutation, 3, compared with the mutations in C5-7. U2 will also be connected with nodes C8-11, and C12-13, which respectively gain new mutation 4 and new mutation 5. It can be seen that without adding U2, connecting C5-7 directly with C8-11 and C12-13 will lead to the gain of new mutation twice on the parallel edges, which is less likely to happen biologically and is now avoided by adding the unobserved node U2 in between the two time points. The resulting tree is shown in **Fig. 1E**.

Once we identify an unobserved node and connect it with the nodes in time points *p* and *p* + 1, we continue with the next round identifying the unobserved nodes from the sorted filtered key-value hash table. But before that, to maintain a tree structure, the nodes in time point *p* + 1 that have been linked with the unobserved node are removed from the hash table because they will no longer be linked with any other unobserved nodes. If this results in any key-value pair whose value has only one node or is empty, we remove the key-value pair as well. This process continues until no more unobserved node can be identified.

It is possible that no node in time point *p* has the mutation set that is a super subset of the mutation set of a key. In this case, we will have to introduce back mutations from a node in time point *p* to the unobserved node. Of all nodes in time point *p*, we will select the node that will lead to the least number of back mutations to be connected with the added unobserved node, i.e., the node that has the least number of mutations that the unobserved node does not have will be selected.

ScLongTree allows more than one layer of unobserved nodes, considering a cascade of branching events may have been missed due to the large gap between two consecutive time points. To construct more than one layer of unobserved nodes, our algorithm will build the structure top-down, from the layer closest to time point *p*, to the layer closest to time point *p* + 1. In more detail, suppose the first layer of unobserved nodes has been identified. These unobserved nodes are linked with the nodes in time point *p* but not time point *p* + 1 because they may be connected to the next layer of unobserved nodes instead of any nodes in time point *p* + 1. In constructing the next layer of unobserved nodes, we treat both the first layer of unobserved nodes and the nodes in time point *p* that have not been linked to any unobserved nodes as the nodes in time point *p*. We repeat the whole process described before. In this case, the nodes in the new layer may have a node in time point *p* as the parent node, or an unobserved node in the previous layer(s) as the parent node. In the case there are more than one node that might be the parent of an unobserved node in the next layer, we give priority to the node in the latest constructed layer. A more detailed explanation accompanied by an example with two layers of unobserved nodes can be found in **Supplemental Section 4**.

### 2.4 Correcting parallel and back mutations

After all iterations of eliminating the potentially spurious subclones, we post-process the inferred tree to correct the parallel and back mutations on the tree using the *k* -Dollo-based constraints [6]. This step does not modify the inferred tree structure or cell clustering, but only refines mutation assignments on the fixed tree. We use the *k* -Dollo-based constraints because parallel mutations are less frequent than back mutations, since most of back mutations occurred due to the copy number losses. Given the inferred tree structure, we first eliminate parallel mutations so that the mutation can occur only once on the tree. We do this by selecting the edge on which the mutation mostly likely occurred, through the calculation of the ratio of the probability when the mutation is present, over the probability when the mutation is absent from a certain branch. Mathematically, suppose a parallel mutation *j* occurred on a set of edges *A*_*j*_. Suppose the descendant nodes of an edge *e* ∈ *A*_*j*_ that have mutation *j* are 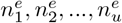. Let 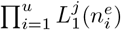 and 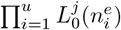 represent the likelihood of having 1 (present) and 0 (absent), respectively, as the inferred genotype for mutation *j* on edge *e*, which is the product of all of its descendant nodes’ likelihood. Such a likelihood can be calculated by **Eq. 3**. For each *e* ∈ *A*_*j*_, we calculate the ratio of these two likelihoods, 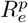, shown in **Eq. 5**. Whichever edge *e* ∈ *A*_*j*_ has the highest 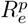 will have mutation *j*. We then remove mutation *j* from other edges in *A*_*j*_, and correspondingly change the inferred genotypes for all descendants of these edges accordingly.

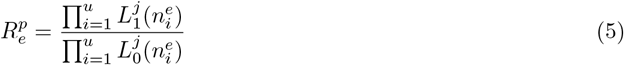

We also added an option to allow parallel mutations, enabling the model to capture cases where the same mutation arises independently on multiple branches, which can occur under selective pressures such as treatment resistance and provides a more biologically realistic representation of tumor evolution.

In the *k* -Dollo model, for a mutation *j*, at maximum *k* edges may lose *j*. Suppose mutation *j* is lost on a set of edges *B*_*j*_. If |*B*_*j*_| ≤ *k*, all back mutations of *j* are accepted. The back mutations can be found by comparing the parent and child nodes’ inferred genotypes, and if the child node does not have a mutation that the parent node has, the mutation is a back mutation. Notice that a mutation may be lost on multiple edges. If a mutation has been found to be lost on more than *k* edges, we perform the following procedures to limit the number of back mutations to *k*. Suppose the descendant nodes of an edge *e* ∈ *B*_*j*_ that have mutation *j* lost are 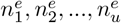. Similar to what we did in removing parallel mutations, let 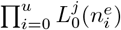 and 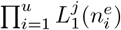 represent the likelihood of having 0 (absent) and 1 (present), respectively, as the inferred genotype for mutation *j* on edge *e*, which is the product of all of its descendant nodes’ likelihood. For each *e* ∈ *B*_*j*_, we calculate the ratio of these two likelihoods, 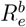, shown in **Eq. 6**. Notice that in contrary to the parallel mutations, the ratio for the back mutations is the likelihood of the “0”s over the likelihood of the “1”s, instead of the likelihood of the “1”s over the likelihood of the “0”s. In other words, our algorithm checks whether it is more likely that the mutation is lost than gained. The top *k* edges that have the highest 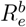 will lose mutation *j*. We then remove the loss of mutation *j* from other edges in *B*_*j*_, and change correspondingly the inferred genotypes for all descendants of these edges. We will re-calculate the final FP and FN values for each time point which are used to eliminate bad BnpC runs described in the next section.

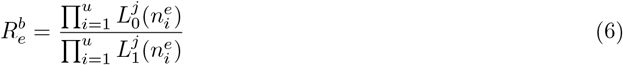

### 2.5 Determining the optimal clustering

We run BnpC for five times since BnpC is non-deterministic. We determined that five is a good number balancing the running time and accuracy after testing on five, ten and fifteen times of BnpC run on the simulated data. More details of this experiment can be found in **Section 3.1**. Each BnpC run will result in a longitudinal subclonal tree. The multiple BnpC runs provide us a chance to eliminate bad BnpC clustering result, as well as the chance to select the best longitudinal subclonal tree resulting from the best BnpC clustering result. To select the best BnpC run and thus the best longitudinal subclonal tree, scLongTree checks the final estimated FP and FN rates for each time point for all five BnpC runs, and eliminates the BnpC runs whose final FP or FN rates are outliers. We define an FP or FN rate as an outlier if it is beyond one standard deviation from the median of the five runs. Of the remaining BnpC runs, we select the run whose final longitudinal subclonal tree has the highest tree support score.

## 3 Results

### 3.1 Simulation

We performed a comprehensive simulation benchmark to compare scLongTree, LACE [21], SCITE [10], SiCloneFit [29] and RobustClone [3]. Of the five methods that we tested, only scLongTree and LACE have been specifically designed for longitudinal scDNA-seq data; the other methods were designed for data collected from one time point. We chose SCITE and SiCloneFit for comparison because both of them have been widely used to infer the phylogenetic tree given one-time scDNA-seq data. The parameter settings of these four methods can be found in **Supplemental Section 6**. In our simulation, we ranged the number of mutations from 50 to 140, and varied the total number of cells among [100, 300, 600, 1000]. The number of mutations and number of unobserved nodes in a tree are controlled by parameters *a* and *µ*, respectively (**Supplemental Table 1**). The number of time points is set as 3 by default.

Here we describe briefly the simulation process. More details of this generative process can be found in **Supplemental Section 2**. We first generate a tree that has multiple time points. To mimic the process when unobserved nodes may occur, we add nodes in between two time points so that they are not represented by any cells sequenced. The number of unobserved nodes is controlled by a variable *u*, which was varied in our simulator to test the robustness of scLongTree. In addition to *u*, we also tested the robustness of scLongTree when different numbers of mutations were involved, which were controlled by the variable *a*. Since sequencing errors are a major compounding factor of placing mutations on a tree, we tested scLongTree when there were varying FP rates, FN rates and missing rates. Finally, error rates may vary at different sequencing time points. We therefore also tested scLongTree when the error rates at each time point varied from each other.

In summary, we varied six variables one at a time to test the robustness of scLongTree compared with other methods, which are FP rate, FN rate, missing rate, the frequency of unobserved nodes *u*, the variance of the error rates among multiple time points, as well as the variable controlling the number of mutations, *a*. For each set of variables, we generated a longitudinal tree via a generative model, and imputed SNVs on the tree. Detailed explanation of the whole generative process as well as the variables used can be found in **Supplemental Section 2**. We list all the variables that we varied in the simulated dataset, along with the values that we used in **Supplemental Table S1**. We evaluated the pairwise SNV accuracy for the five methods based on the trees they inferred. Pairwise SNV accuracy evaluates the percentage of the correctly inferred pairwise SNV relationship among all pairs of SNVs. Two SNVs may have three possible pairwise relationships: ancestral, same edge, and parallel. A detailed explanation of how the pairwise SNV accuracy is calculated is provided in **Supplemental Section 8.1**. We show the pairwise SNV accuracy for the six variables from scLongTree, LACE, SCITE, SiCloneFit and RobustClone in **Fig. 2**. In all cases except one, scLongTree outperformed all other methods. The only exception was when the false positive rate reached 0.05, where LACE performed slightly better (**Fig. 2a**). However, scLongTree demonstrated robustness across a varying range of error rates (**Fig. 2e**). Notably, when the mutation number is high, LACE fails to complete within 24 hours, as observed when *a* = 5.1 (average mutation number around 140) in **Fig. 2f**. Additionally, due to the design of scLongTree that allows unobserved nodes, scLongTree’s pairwise SNV accuracy remained consistently close to 1 for all degrees of unobserved nodes (**Fig. 2d**), whereas the other methods showed pairwise SNV accuracy lower than and less robust than scLongTree.

**Figure 2:**
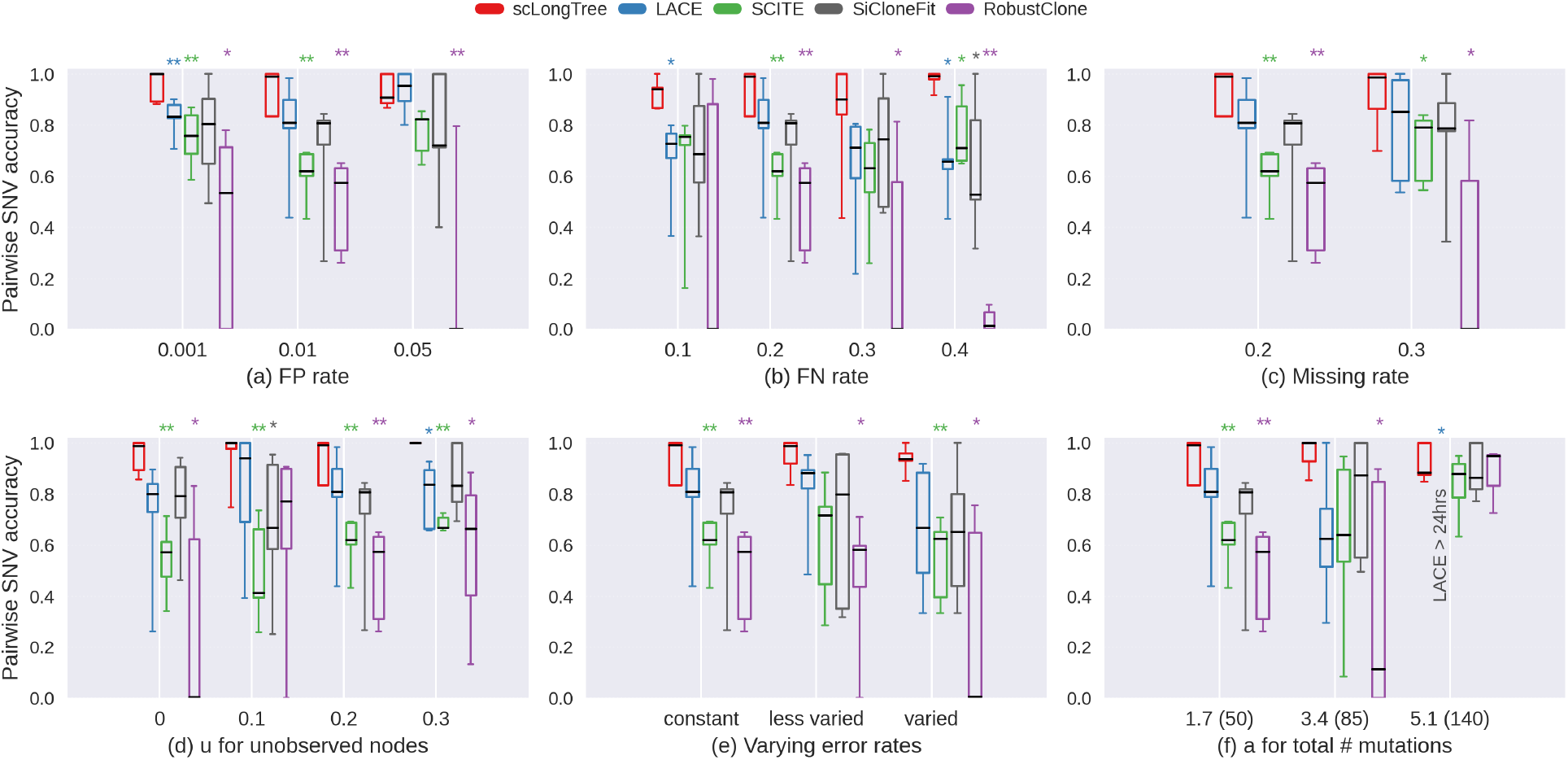
Boxplots are shown for pairwise SNV accuracy for scLongTree (red), LACE (blue), SCITE (green), SiCloneFit (black) and RobustClone (purple). Six variables were varied to test the robustness of these five methods, including FP rate (a), FN rate (b), missing rate (c), variable *u* that controls the number of unobserved node (d), the range of error rate across different time points (e), and variable *a* (f) that controls the number of mutations. The average numbers of mutations in (f) corresponding to a certain value of *a* are inside the parenthesis next to the values of *a*. Median of each boxplot was highlighted with black horizontal line. Each set of variables was repeated for five times to avoid extreme cases. Statistical significance of the Student t-test between the competing method and scLongTree is indicated by asterisks: * (p *<* 0.05), and ** (p *<* 0.01).

As a complementary measuring metric, we also measured the Robinson-Foulds (RF) accuracy between the inferred tree and the ground truth tree, the boxplots of which were in **Supplemental Fig. S7**. Overall, scLongTree demonstrates superior performance compared with all other methods. Notably, its advantage becomes more evident as the number of mutations grows, as reflected by its higher RF accuracy to the ground-truth tree in high-mutation settings. We noted that RobustClone consistently underperformed other methods, reaching zero V-measure for quite a few variables shown in **Fig. 2** and zero RF accuracies for all variables shown in **Supplemental Fig. S7**. We therefore eliminated RobustClone for the following analysis. In addition, we evaluated scLongTree’s imputation accuracy for missing entries (**Supplemental Fig. S8**). To assess imputation accuracy, we first matched each inferred node to its corresponding ground truth node by calculating the Jaccard index between all pairs of nodes and selecting the ground truth node with the highest Jaccard index at the same time point for each inferred node. For each cell-mutation pair with a missing value in the original data, we then compared the imputed genotype (derived from the consensus genotype of the node containing the cell) to the true genotype from the corresponding ground truth node. Imputation accuracy was calculated as the percentage of correctly imputed mutation genotypes across all missing values. The median accuracy was generally above 0.7, indicating that scLongTree effectively imputed missing data.

In addition to the tree accuracy, we also investigated scLongTree’s capability of improving the clustering accuracy. For this experiment, we tested the robustness of scLongTree’s clustering accuracy with respect to the number of BnpC runs, varying this parameter to 5 (default), 10, and 15 runs. For comparison, we also included the BnpC result with the highest likelihood (see **Supplemental Section 7**) to examine whether scLongTree could achieve improved clustering when incorporating tree information. Furthermore, we compared against the state-of-the-art clustering method SCG [23]. **Supplemental Figure S9** presents boxplots of the V-measures for all methods across the different conditions in the simulated data. Throughout all variables, scLongTree’s V-measure stayed high near 1 for 5, 10 and 15 BnpC runs, showing that more BnpC runs do not necessarily increase scLongTree’s accuracy. Thus five BnpC runs is an appropriate number balancing accuracy and runtime. The BnpC run with the highest likelihood performs second best, while SCG exhibits the lowest V-measure. This outcome is expected, as scLongTree leverages the inferred tree structure to refine clustering, an advantage not available to BnpC. BnpC and SCG are otherwise comparable cell clustering tools. Consistent with what was observed in [11], BnpC’s clustering accuracy generally outperforms SCG. Selecting the BnpC run with the highest likelihood further improves its V-measure, though it still falls short of scLongTree. These results demonstrate that incorporating tree information is essential for refining cell clustering and achieving optimal clustering accuracy.

We then further evaluated the accuracy of SNV placement in terms of time points. For each method, we calculated the recall as the percentage of the SNVs correctly placed on a time point over all SNVs placed at the same time point in the ground truth data, and precision as the percentage of the SNVs correctly placed on a time point over the total SNVs placed at the same time point by the method. The SNV placement accuracy is defined as the harmonic mean of the average recall and average precision over all time points. More details of this evaluation metric can be found in **Supplemental Section 8.2**. Since SCITE and SiCloneFit do not utilize longitudinal information and therefore do not assign mutations to specific time points, we infer the time point of each mutation based on the time points of the cells that carry it. Details can be found in **Supplemental Sections 9**. We found that scLongTree is superior to LACE in terms of the SNV placement accuracy (**Fig. 3**). ScLongTree’s median accuracy stayed near 1 and remained highest among all four methods. LACE’s SNV placement accuracy is lower than scLongTree, but higher than SCITE and SiCloneFit, showing the advantage of considering the time points of the cells since SCITE and SiCloneFit do not consider the time points. Nevertheless, among the two methods that take as the input a longitudinal tree, scLongTree outperforms LACE in terms of SNV placement accuracy. It is also worth to note that LACE cannot produce the result when the number of mutations is high above 100 (**Fig. 3f**) within 24 hours.

**Figure 3:**
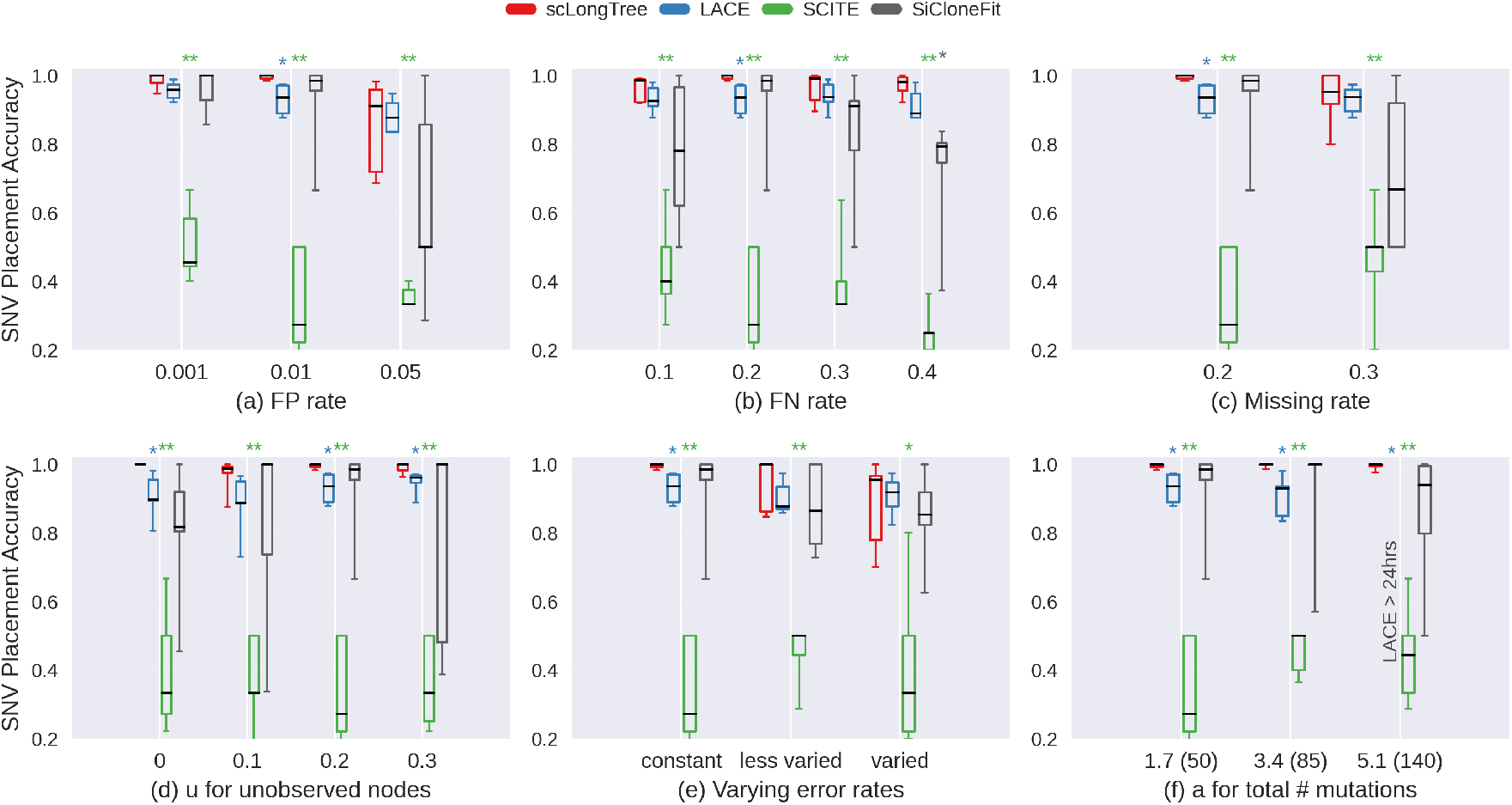
Boxplots are shown for SNV placement accuracy for scLongTree (red), LACE (blue), SCITE (green), SiCloneFit (black). Six variables were varied to test the robustness of these four methods, including FP rate (a), FN rate (b), missing rate (c), variable *u* that controls the number of unobserved node (d), the range of error rate across different time points (e), and variable *a* (f) that controls the number of mutations. The average numbers of mutations in (f) corresponding to a certain value of *a* are inside the parenthesis next to the values of *a*. Median of each boxplot was highlighted with black horizontal line. Each set of variables were repeated for five times to avoid extreme cases. Statistical significance of the Student t-test is indicated by asterisks: * (p *<* 0.05), and ** (p *<* 0.01).

To further investigate the relationship between pairwise SNV accuracy and phylogenetic distance, we compared accuracy for parent-child versus grandparent-grandchild relationships for scLongTree (**Supplemental Fig. S10**). Consistent with observations in bulk sequencing where pairwise accuracy decreases as trees grow larger, we found that pairwise SNV accuracy declined as the distance between node pairs increased. This pattern of grandparent-grandchild pairs showing lower accuracy than direct parent-child pairs reflects the greater phylogenetic distance between more distantly related nodes.

To assess scLongTree’s ability to infer unobserved nodes, the internal nodes in the phylogeny that lack sampled cells, we analyzed the detection accuracy, the F1 score, across all simulated datasets. We matched inferred unobserved nodes to ground truth unobserved nodes by requiring that their observed parent or daughter nodes share a Jaccard Index ≥ 0.75 based on the cells they contain, or that both unobserved nodes are the daughter nodes of the root, whose node ID is shared between ground truth and inferred trees. Our analysis revealed that scLongTree achieved an average of 1.73 true positive and 0.59 false positive unobserved nodes per dataset (median: 2 and 0, respectively). The size distribution of inferred unobserved nodes (range: 2-6 cells, average: 2.31, median: 2) closely matched that of ground truth unobserved nodes (range: 2-6 cells, average: 2.46, median: 2). The F1 scores for unobserved node inference across different simulation conditions are shown in **Supplemental Fig. S11**, demonstrating scLongTree’s ability to recover internal phylogenetic structure even when intermediate lineages are not directly sampled. To evaluate whether inferring unobserved nodes improves phylogenetic accuracy, we compared the RF accuracy between inferred and ground truth trees with and without unobserved nodes included (**Supplemental Fig. S12**). In the latter case, we directly connected the offspring of each unobserved node to its parent, effectively collapsing the unobserved internal nodes. Trees that retained inferred unobserved nodes exhibited higher RF accuracy in the majority of simulated datasets, demonstrating that recovering these internal nodes improves the overall accuracy of the phylogenetic reconstruction.

Considering BnpC’s non-deterministic nature, we further examined BnpC regarding its stability since BnpC provides the initial clustering result to scLongTree. **Supplemental Fig. S13** showed the V-measure of BnpC using each of the BnpC run as the ground truth for varying FP and FN rates, each variable repeated for five times generating five trees (rep1-rep5). BnpC’s performance is relatively stable in multiple runs. The average run-to-run similarity with the V-measure as the metric stays near 1 when the error rate is small (FP = 0.001, FN = 0.1). The stability decreases with the increase of the FP rate (**Supplemental Fig. S13a**) and FN rate (**Supplemental Fig. S13b**). Nevertheless, most of the V-measure measuring run-to-run similarity stays above 0.8, showing the stability of BnpC. In addition, we tested the scLongTree’s clustering accuracy by running BnpC pooling and not pooling cells across time points together, respectively (**Supplemental Fig. S14**), which showed that difference of the V-measure is negligible between these two settings.

We then looked into the running time comparing scLongTree, LACE, SCITE and SiCloneFit with increasing number of SNVs (**Fig. 4**). Of the four methods, SCITE’s running time remained the lowest. LACE’s running time did not increase when the number of SNVs increased from about 50 to 85. However, when the number of SNVs increased to 140, the running time of LACE increased dramatically, to more than 24 hours. ScLongTree’s running time was the highest in these simulations, except when the SNV number was high. This is because the reported computation time includes running BnpC for five times, and the majority of the runnning time was spent on running BnpC. It typically took less than 10 seconds for scLongTree to come up with the final tree after BnpC results were obtained, and thus its own running time was negligible compared with the time spent on running BnpC. **Supplemental Fig. S15** includes the running time of scLongTree without considering BnpC runs. We list the median running time of all four methods in **Supplemental Table S2**, which also contains the running time of scLongTree with and without the BnpC runs.

**Figure 4:**
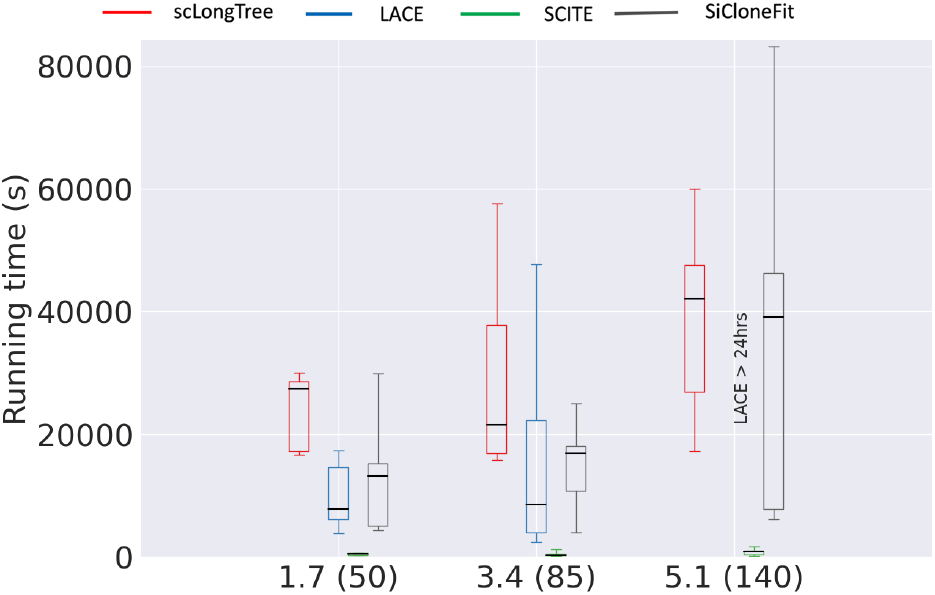
Boxplots are shown for running time when we varied variable *a* for the trees inferred from scLongTree (red), LACE (blue), SCITE (green) and SiCloneFit (black). The corresponding average number of mutations are inside the parenthesis next to the values of *a*. Median of each boxplot was highlighted in black horizontal line.

In all, scLongTree has been shown to be advantageous over other methods in the simulated dataset.

### 3.2 Real data

#### Validation on SA501 dataset

We applied scLongTree to a real triple-negative breast tumor sample, SA501, for which longitudinal targeted patient-derived xenograft scDNA-seq data were collected at three time points, *X*1, *X*2, and *X*4 [5]. A total of 27, 36 and 27 cells were sequenced, at each time point, respectively. LACE [21] was also applied to SA501, in which 20 out of the 55 variants identified in [5] were selected to construct the longitudinal tree.

We first applied scLongTree to the same 20 variants selected by LACE to infer the longitudinal subclonal tree, as shown in **Fig. 5a**. We show the corresponding heatmap of our inferred genotypes in **Fig. 5b**. We found that our inferred tree was the same as both LACE’s result and the tree inferred in the original paper that published SA501’s data [5]. The latter was further confirmed with biological means in the same study [5]. Just like LACE, we numbered these 20 variants from 0 to 19. According to the tree that scLongTree, LACE, and the original study [5] inferred, variants 0-6 occurred at the trunk. The tree then branched out. One branch obtained variants 11-13, and the other branch obtained variants 7-10. On the first branch, variants 14-18 were further obtained after variants 11-13. These were all shown in the time point X1. X2 was mainly the same as X1, but variant 19 was gained after X2 and was shown in time point X4. In the trees in **Fig. 5**, we used dark pink, yellow, and purple to color the nodes in time points X1, X2 and X4, respectively.

**Figure 5:**
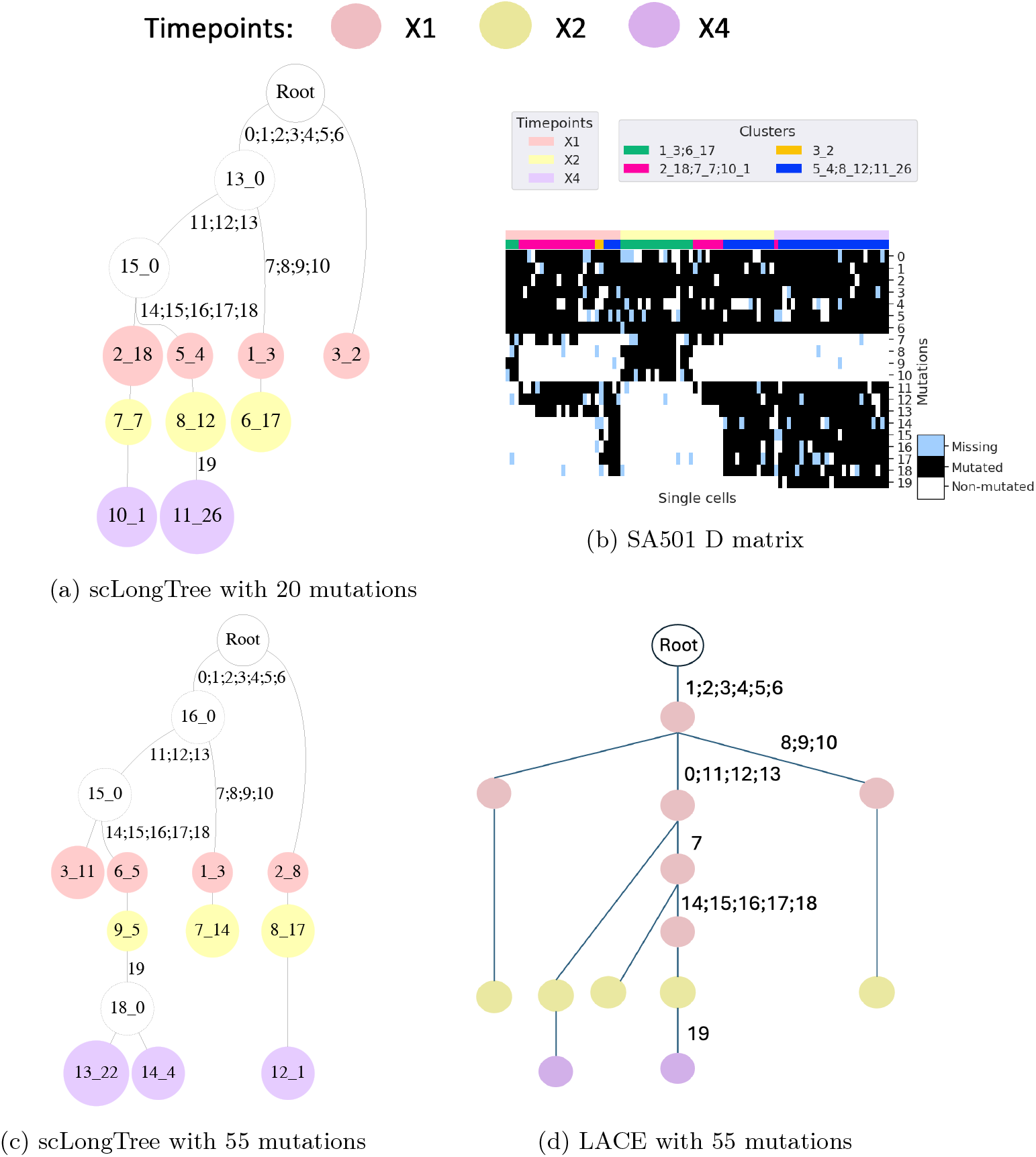
Sample SA501: (a) scLongTree’s inferred longitudinal subclonal tree for sample SA501 with 20 mutations. Annotated on the edges are the new mutations. The nodes are named by the node number followed by the number of cells, the two of which are separated by “ “. For example, node “2 18” is node 2 and it has 18 cells. The nodes with dotted circles are unobserved nodes which do not have any cells sequenced. (b) Corresponding SA501 heatmap matrix for 20 mutations. The horizontal color bars indicate time points and clusters of the cells, respectively. The entries have blue, black and white colors representing missing entries, 1 and 0 entries, respectively, as observed from the data. (c) scLongTree’s inferred longitudinal subclonal tree for sample SA501 with 55 mutations where only 20 mutations are presented to show that it is consistent with (a). **Supplemental Fig. S16** shows the longitudinal tree with all 55 mutations. (d) LACE’s inferred longitudinal subclonal tree for SA501 with 55 mutations where only 20 mutations are marked for comparison.

We then further checked whether we could infer the same tree structure using not only the 20 variants that LACE used, but also the rest of the 35 variants. We call the dataset with the 20 variants the “small SA501” dataset and the dataset with all 55 variants the “large SA501” dataset. Our assumption is that the tree structure inferred using the small SA501 dataset and the large SA501 dataset should be consistent. In other words, adding the 35 variants should not violate the tree structure that was inferred using the 20 variants if we do not consider the extra branches resulting from the extra 35 variants. We thus applied scLongTree to the 55 variants and inferred the longitudinal subclonal tree as shown in **Fig. 5c**, which shows only the 20 variants for the comparison with the tree inferred using only the small SA501 dataset. The original tree inferred from the large SA501 dataset that shows the annotation of all 55 variants is in **Supplemental Fig. S16**, in which the black numbers on the black edges are from the 55 variants, and blue numbers on the blue edges are the corresponding indices of the 20 variants. Comparing the scLongTree inferred trees given the small SA501 dataset in **Fig. 5a** and the large SA501 dataset in **Fig. 5c**, we found the two tree structures had exactly the same pairwise relationship of the variants, showing the robustness of scLongTree to the number of variants given. We reasoned that the tree topology using the 55 mutations does not differ from that using 20 mutations because the additional 35 mutations have higher missing rates, limiting their influence on clustering and tree construction relative to the 20 high-confidence mutations. We also noticed that mutation 21 in the 55 mutations uniquely linked an additional unobserved node 18 0 with node 13 22 (**Supplemental Fig. S16**). This mutation lies within the pseudogene CDRT15P3 and is not known to play a functional role in breast cancer. Its late, subclonal occurrence is consistent with a passenger mutation. Nevertheless, such mutations are informative for resolving lineage relationships and temporal ordering, highlighting the value of unobserved nodes in accurately representing tumor evolutionary history.

We then applied LACE to the large SA501 dataset to infer the longitudinal subclone tree shown in **Fig. 5d**. For comparison, in **Fig. 5d**, we only showed the small SA501 dataset based on the tree structure inferred from LACE given the large SA501 dataset. We found that LACE’s inferred tree given the large SA501 dataset differed from its inferred tree given the small SA501 in terms of the pairwise relationship of the variants. Specifically, variant 7 should have been placed on the branch of 8-10, but LACE’s inferred tree given the large SA501 dataset placed 7 on the other branch, underneath variants 11-13. In addition, variant 0 should have been placed on the trunk before the branching happened. However, in the tree inferred by LACE from the large SA501 dataset, variant 0 was incorrectly placed on one of the branches, together with variants 11-13, leaving the other branch containing variants 8-10 without variant 0. This comparison showed that LACE’s results are not robust to the number of variants used to infer the tree. In contrast, scLongTree showed consistency of the tree structures given the large and small SA501 datasets.

### Validation on a larger AML dataset

To assess the scalability and generalizability of scLongTree beyond SA501, we applied our method to the longitudinal AML107 dataset from Morita et al. [18], which contains substantially more cells (2,817 cells at diagnosis and 1,800 cells at relapse). This dataset represents a typical AML case with longitudinal single-cell DNA sequencing data collected at two time points during disease progression. Using scLongTree, we successfully reconstructed the subclonal evolutionary tree and identified the temporal ordering of driver mutations. Consistent with the original publication, our analysis revealed that TP53 mutation occurred as an early event, followed by acquisition of DNMT3A mutation in the evolved subclone. The inferred tree topology (shown in **Supplemental Fig. S17**) was highly concordant with the phylogenetic tree reported in the original study, demonstrating that scLongTree maintains accuracy when scaling to datasets with 46 times more cells than SA501. This validation confirms the robustness of our method across different sample sizes and AML genetic contexts, supporting its utility for analyzing longitudinal single-cell sequencing data in diverse clinical scenarios.

## 4 Discussion

ScLongTree is a computational tool that can infer the phylogenetic tree based on longitudinal scDNA-seq data. Multiple approaches have been integrated into scLongTree to obtain a high accuracy, including clustering cells in each time point to accurately infer error rates in each time point, using statistical analysis to eliminate small spurious subclones, and having a rigorous control of the FP, FN and missing rates at each time point to avoid inflated false positives and false negatives. In addition, we developed an efficient algorithm to construct the phylogenetic tree, which can infer unobserved subclones that occur in between two consecutive time points and thus are not represented by any cells. In simulation, we compared scLongTree with the existing method LACE that infers subclonal trees from longitudinal scDNA-seq data, as well as three state-of-the-art methods that infer phylogenetic trees based on scDNA-seq data from one time point, SCITE, SiCloneFit and RobustClone. We found that scLongTree outperformed all other three methods for all six experiments conducted in terms of pairwise SNV accuracy. In addition, scLongTree was scalable to hundreds of mutations, whereas LACE could not finish the run within 24 hours when there were over 140 mutations in the input matrix. Application of scLongTree to the real, triple-negative breast cancer dataset SA501 showed that not only was scLongTree able to recover the evolutionary history of the cancer, but also was able to maintain the consistency of the pairwise relationship of the mutations when using a larger number of mutations, whereas LACE failed to show such a consistency. This shows that scLongTree is more robust to the number of mutations in inferring the longitudinal subclonal trees. Furthermore, validation on the AML107 dataset, which contains 4,600 cells, demonstrated that scLongTree is robust to the number of cells.

Future directions include combining the clustering and tree inference in one step, which although may increase the computational complexity, may result in a higher accuracy of the inferred tree. This is because clustering can benefit from the knowledge of the tree structure, and thus inferring clustering and the tree structure simultaneously is more advantageous in terms of accuracy. In this context, incorporating explicit evolutionary constraints, such as the *k* -Dollo model, directly into joint clustering and tree inference may further improve the biological plausibility of the inferred topology.

Another interesting direction is to decrease the false negative rate by inferring the allele dropout (ADO) regions, which becomes possible with the advancement of the understanding of the single cell whole genome amplification error profiles [32]. Given decreased false negative rate in the observed genotypes, the inferred longitudinal subclonal tree will be more accurate. It is also possible to probabilistically score each FN that may be due to the ADO based on both the position of the mutation and the inferred longitudinal tree. In this way we can decrease the FN rate given not only the genomic positions of the FNs, but also an inferred longitudinal subclonal tree structure. The decreased FN rate can further improve the tree accuracy in an iterative manner.

A potential limitation of scLongTree is that it links subclones only between consecutive time points. Consequently, if a subclone is observed at time point *p* and *p* + 2 but not at *p* + 1, the current algorithm does not directly connect these nodes. This design reflects a conservative approach, prioritizing connections supported by observed intermediate states and reducing the risk of overinterpreting gaps caused by sparse sampling or spatial heterogeneity. Extending the algorithm to infer links across non-consecutive time points could be a useful direction for future development, particularly in studies with sparse temporal sampling.

In contrast to scLongTree, finite-sites models such as SiFit [27] do not constrain the number of times a site can change state. They typically model mutational dynamics as a continuous-time Markov chain over genotype states, with separate rate parameters for gains, losses, and LOH events. This means every site can in principle undergo any sequence of state transitions, and the likelihood of the observed data is typically computed by marginalizing over all such histories on a given tree. In scenarios with pervasive convergent evolution driven by strong positive selection where the same mutation independently arises in distinct lineages or in tumor contexts with widespread copy number driven mutation loss that affects many sites simultaneously, this richer parameterization may yield more accurate phylogenetic inference by assigning explicit probabilities to each possible evolutionary path rather than applying a fixed bound on back-mutation counts. However, this expressiveness comes with important limitations that motivated scLongTree’s design choices. First is scalability. Finite-site models such as SiFit rely on MCMC to explore the joint space of tree topologies and error rates, which becomes computationally prohibitive as the number of cells and mutations grows. scLongTree is designed to scale to hundreds of mutations and thousands of cells, a regime where MCMC-based finite-sites inference is not practical. Second, is identifiability. With a fully reversible mutation model, the number of rate parameters to be estimated grows with the number of sites and the complexity of the transition model. In datasets with moderate cell counts which are typical of longitudinal scDNA-seq studies, this can lead to poorly constrained parameter estimates and overfitting. In contrast, the *k* -Dollo constraint in scLongTree acts as a biologically motivated regularizer to avoid overfitting. Third, and critically for scLongTree’s setting, finite-sites models such as SiFit do not exploit longitudinal information. They treat all cells as arising from the same time point, and therefore cannot leverage the temporal ordering of subclones to disambiguate whether an apparent mutation loss is a true biological event or an artifact of sparse sampling at a given time point, a disambiguation that scLongTree’s time-point-aware tree construction directly supports. We acknowledge that scLongTree’s heuristic approach may be less well-suited to datasets where convergent evolution is pervasive or where copy number driven loss affects a substantial fraction of sites in a manner not captured by the *k* parameter. Nevertheless, scLongTree is advantageous in scalability, identifiability and exploiting longitudinal information compared with finite-sites models. In summary, scLongTree offers a practical and accurate balance between biological realism, scalability, and the ability to exploit multi-time-point structure. Future work includes exploring hybrid approaches that apply a more probabilistic treatment of mutation loss specifically at sites flagged as likely copy number variation affected, while retaining the combinatorial efficiency of scLongTree’s tree construction for the remainder.

Lastly, scLongTree’s flexibility in inferring multiple layers of unobserved nodes and allowing back mutations may lead to overfitting issues, which could produce complex trees in datasets with limited or noisy observations. While we have not observed any overfitting in the simulated and real data, and the current design already incorporated implicit regularization through constraints on the depth of unobserved nodes, data-supported node construction, and the *k* -Dollo limit on back mutations, adding an explicit regularization term could further penalize overly complex tree structures. Such a term, for example based on Bayesian priors or information-theoretic criteria, could provide a principled approach to balance model flexibility with parsimony, and represents a promising direction for future development.

## 5 Data availability

Triple-negative breast tumor sample SA501 was obtained from the European Genome-phenome Archive under accession number EGAS00001000952. Further processed data was available in http://www.cbioportal.org/patient/clinicalData?studyId=brca_bccrc_xenograft_2014&caseId=SA501. ScLongTree is freely available on https://github.com/compbio-mallory/sc_longitudinal_infer.

## 6 Acknowledgement

We would like to thank Daniele Ramazzotti for helping us interpret LACE’s result. We also would like to thank Adi Steif and Peter Eirew for sharing the SA501 raw tables for our analysis.

