## Supplemental Text for "scLongTree: an accurate computational tool to infer the longitudinal tree for single-cell DNA sequencing data"

### Supplemental Methods of "scLongTree: an accurate computational tool to infer the longitudinal tree for scDNAseq data"

#### Contents

**1 Advantages of having time points for interpreting the biology. 3**

**2 Description of our simulator 4**

**3 Determining the threshold for eliminating clusters 8**

**4 Illustration of Tree Inference Algorithm 8**

**5 Reassigning cells to new clusters 13**

**6 Running scLongTree, LACE, SiCloneFit, and SCITE for evaluation 13**

**7 Selecting the best BnpC run 14**

**8 Evaluation metrics 15**

**9 Determining SCITE and SiCloneFit's SNV placement accuracy 16**

*These authors contributed equally to this work.
†These authors contributed equally to this work.

**10 Supplemental Tables** **17**

**11 Supplemental Figures** **18**

**12 Supplemental Algorithm S1** **27**

### 1 Advantages of having time points for interpreting the biology.

The only extra signal used in the longitudinal data compared with the one time-point data is the time point, i.e., each cell has not only its observed genotype, but also the time point when it was sequenced.

A longitudinal subclonal tree is different from a traditional phylogeny in that it takes into account the time point a cell is sequenced. Such time points can constrain the tree better, because the cells have to stay on their own time point on the tree, whereas the tree still has to follow evolution according to certain error rates.

It is possible to infer the phylogeny without considering the time points using traditional methods such as SCITE, SiFit, SiCloneFit, etc, followed by labeling the cells with their time points. However, this is not preferred and one obvious reason is that the tree algorithms without considering the time points of the cells do not necessarily respect the time points. For example, cells from the same time point may be very far away from each other in a hierarchy. Cells sequenced at a later stage may be placed closer to the root, inconsistent with the time points. Here we list a few advantages of considering time points while inferring the phylogenetic tree with illustrations. First, the time points help to resolve the mutation order, especially when the parallel or back mutations are involved. For example, as shown in the left panel of **Supplemental Fig. S1**, disregard the sequencing errors, suppose 100% cells from T1 has mutations A, B, 50% cells from T2 have mutations A, B, C, and the rest of the 50% cells from T2 have mutations A, D. Without the time points, a perfect phylogeny can be inferred, which has A for all cells, then branches out to B and D, in which B is further followed by C, seen in the right panel in **Supplemental Fig. S1**. However, since 100% cells from T1 have A and B, if the sampling of the cells covers the entire biological area, the data strongly indicates that A and B are co-existing instead of A preceding B. Moreover, the loss of B cannot be detected in the case of not considering the time points.

Second, incorporating time points helps to infer the unobserved nodes. For example, as shown in the left panel of **Supplemental Fig. S2**, suppose 100% cells from T1 have mutations A, B, 50% cells from T2 have mutations A, C, E, and the rest of the 50% cells from T2 have mutations A, D, E. According to scLongTree's tree algorithm, an unobserved node that has mutations A, E will be inferred. Thus this unobserved node gained a new mutation E and lost mutation B between T1 and T2. Without the time points, the tree on the right panel of **Supplemental Fig. S2** will be inferred, which indicates that B and E are gained in parallel, and A precedes B, inconsistent with the observation that 100% cell in T1 had A, B and thus A and B coexist. It also indicates that there are two unobserved nodes, which have mutations A, and A, E respectively. However, the inferred unobserved node with mutation A has never existed. It also failed to infer the back mutation of B, unlike what a longitudinal subclonal

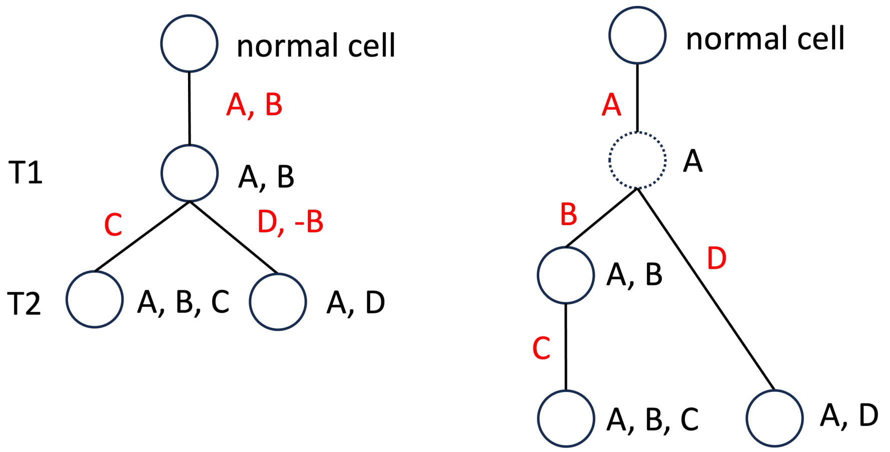


Supplemental Fig. S1: Trees inferred by considering the time point (left panel), and not considering the time point (right panel), emphasizing the longitudinal subclone tree's ability to resolve mutation order. Dotted circles are the unobserved nodes.

tree would have inferred.

Third, incorporating time points helps to infer clonal dynamics. For example, as shown in the left panel of **Supplemental Fig. S3**, suppose 50% cells from T1 have mutations {A}, 50% cells from the same time point have mutations {A, B}. All cells in T2 have mutations {A, C}. ScLongTree will infer the tree to have obtained A, followed by B, and the subclone with mutations {A, B} on T1 disappeared in T2, perhaps from therapies or selective pressure. However, without the time point, a perfect phylogeny will infer that mutation A is followed by B and C in parallel, not indicating the loss of the subclone with mutation {A, B} shown in the right panel of **Supplemental Fig. S3**.

#### 2 Description of our simulator

The simulation is a generative process that simulates a tree. On the tree, the nodes represent subclones, and the edges represent a relationship between the parent and the child. In addition to the mutations that the parent has, a child may have new mutations. We pre-define the total number of time points to be 3, whereas the first two time points happened 15 and 7 years ago, and the third time point is current (i.e., 0 years ago).

We then uniformly sample the total number of cells sequenced at each time point from [100, 300, 600, 1000], like what LACE did [3].

On the tree, each node has an interval which is within the range of 0 and 1. At any time, all the leaf nodes' intervals add up to 1. The purpose of having

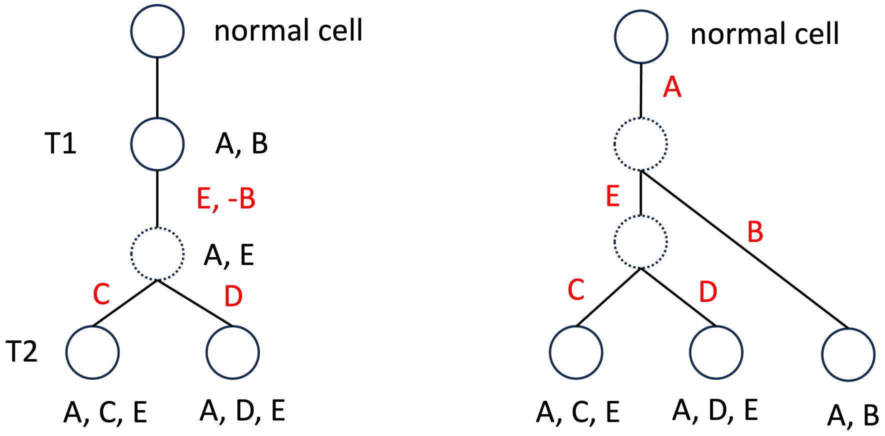


Supplemental Fig. S2: Trees inferred by considering the time point (left panel), and not considering the time point (right panel), emphasizing the longitudinal subclone tree's ability to infer unobserved nodes. Dotted circles are the unobserved nodes.

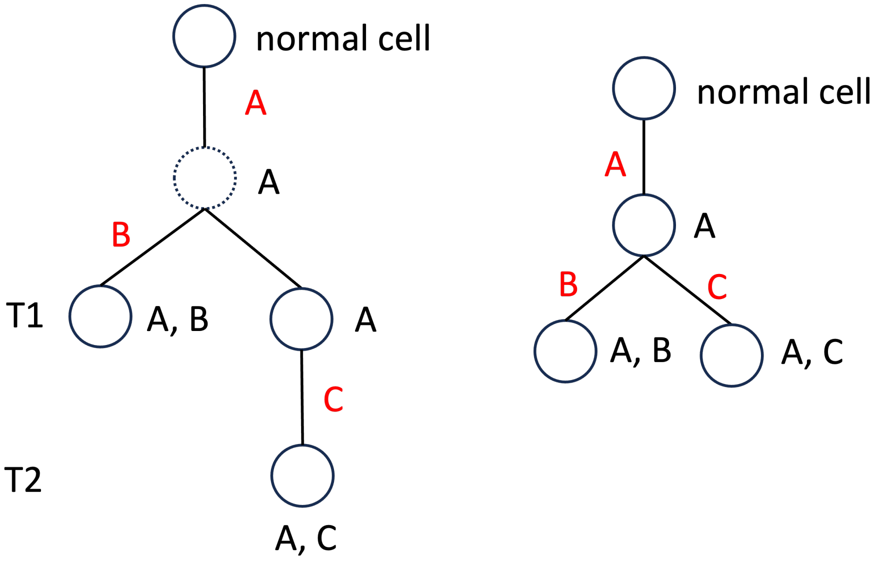


Supplemental Fig. S3: Trees inferred by considering the time point (left panel), and not considering the time point (right panel), emphasizing the longitudinal subclone tree's ability to infer clonal dynamics. Dotted circles are the unobserved nodes.

the intervals for each node is for distributing the cells. At each time point, the percentage of the cells a node may be distributed is linear to the interval the node has. The tree generation process starts with a root node which represents a normal clone without any mutations and is on time point 1. The root node's interval is [0, 1]. In the very beginning, this root node is a leaf node since at this moment, we have not added any child nodes to it. We then start the generative process and add nodes to an existing leaf node. We do this iteratively. At every iteration, we select a leaf node. For the selected leaf node, we select one of the three options, 1) the leaf node splits into two child nodes; 2) the leaf has only one child node, and the child node inherit the same set of mutations as this selected node and does not have any new mutation; and 3) the leaf node is deemed to not have any child node. We select these three options with probability of $1 - v2$, $v2 - v1$ and $v1$, respectively. Here $v1$ and $v2$ are the two numbers between 0 and 1, and $v2 > v1$. $v1$ and $v2$ take part in controlling the tree structure.

We then further decide whether the child nodes shall have new mutations compared with the parent node if the parent node has been decided to split into two child nodes. Specifically, both child nodes are subject to have new mutations with the probability of $\theta$. Otherwise one of the child nodes will not have new mutations. Here $\theta$ is a variable that is tunable in the simulator.

This whole process of having zero, one persistent or two child nodes, whereas the child nodes may or may not have new mutations mimics the real case scenario of subclonal growth. If a node is decided to have at least one child node, the child nodes become leaf nodes and the parent node is no longer a leaf node. The union of the child nodes' intervals is the same as the parent node's interval. If a node is decided to have two child nodes, the two child nodes split the interval the parent node has, and the two child nodes' interval ratio is decided by a sample from the beta distribution whose $\alpha = 0.5$, $\beta = 0.2$. In this way, we control the size of the subclones by controlling the parameters of the beta distribution. Such a usage of Beta splitting model can also be found in [1, 2, 4].

To decide the child nodes' time point, we consider whether the child nodes shall be at the next sequencing time point, or in between the current and next sequencing time point and thus represent an unobserved subclone. We consider unobserved subclones in our simulation because in the real case scenario, it is possible that some subclones with a unique set of mutations occur in between two sequencing time points, whereas such subclones are not represented by any cells being sequenced. In more detail, at each time point, we randomly select a node and decide that this node has two child nodes that will be unobserved subclones by probability $u$. $u$ can be tuned in the simulator. To increase the complexity of the tree, root node always has two child nodes that are unobserved subclones. Notice that a node selected to split into two child nodes representing unobserved subclones is not subject to the probabilities of not having two child nodes such as $v2 - v1$ and $v1$. This process of selecting a leaf node, deciding whether it will split, or is inherited by one node, or does not have any child nodes, stops when all the nodes except those on the last time point have been selected and settled.

Once the tree structure is decided, next, we place the mutations on the edges of the tree based on the branch length, whereas the branch length is the number of years in between two consecutive sequencing time points. To decide the branch lengths that connect the unobserved nodes, we determine the time points of the unobserved nodes in between two consecutive sequencing time points $t1$ and $t2$ by sampling from a Beta distribution whereas $\alpha = \beta = 2$. Suppose such a sample is $s$. The time point of the unobserved node is $t1 + (t2 - t1)s$. Finally, given the branch length of each branch, say $b$, on the tree, the number of mutations each branch is distributed is $ab$, in which $a$ is a new parameter that we varied to test the robustness of scLongTree. $a$ represents the mutation rate. Here we varied $a$ from 1.7, to 3.4, to 5.1, which would result in the total mutation numbers to be around 50, 85 and 140.

Then we distribute the cells on the observed nodes. First, we normalize the intervals of all the observed nodes at each sequencing time point. The normalization is necessary since some nodes do not have any child nodes, and thus shrink the sum of the interval lengths and make it smaller than 1. We then distribute the number of cells according to the length of the interval whereas the total number of cells of each time point has been predefined described above. Notice that since the unobserved nodes do not have any cell sequenced, we do not distribute any cells on the unobserved nodes. Thus all cells are distributed on the nodes that are placed on the sequenced time points.

After the cells are placed on the tree, the $G$ matrix is decided. A cell's underlying true genotype is decided by the mutations on the path from root to the cell's node. For each cell, we then decide the observed data $D$ which has false positive (FP), false negative (FN) and missing entries. Specifically, given the FP rate and FN rate, we flip from 0 to 1 according to the FP rate, and from 1 to 0 according to the FN rate. We finally flip from 0 and 1 to 3 according to the missing rate. In our simulation, FP, FN and missing rates were the three variables that we varied to test the robustness of scLongTree, respectively. We selected FP rate from 0.001, 0.01, 0.03, and 0.05, and FN rate from 0.1, 0.2, 0.3 and 0.4. Missing rate was selected from 0.2 and 0.3. These numbers were selected based on our prior knowledge of the error rates [1, 3].

We performed the simulation study given different FP, FN and missing rates in two different ways. In the first way, we sampled these error rates once and applied the sampled error rates to all time points. Thus the error rates remained the same across different time points, and thus is called "constant". In the second way, we allowed different error rates at different sequencing time points. Specifically, we tested scLongTree in two different modes in the second way. The first mode was called "less varied", in which the FP rate was sampled between 0.01 and 0.03, and FN rate was sampled between 0.2 and 0.3, whereas missing rate was fixed at 0.2. Each of the sampled error rate was applied to one time point, and thus the FP and FN rates might be different at different time points. The second mode was called "more varied", in which all the error rates were sampled from all ranges tested, i.e., FP rate was sampled from 0.001, 0.01, 0.03 and 0.05; FN rate was sampled from 0.1, 0.2, 0.3 and 0.4, and missing rate was sampled from 0.2 and 0.3. Like the "less varied", "more varied" mode also sampled the error rate for each time point and thus the error rates at different time points might be different. This simulation was to test scLongTree's robustness when the error rates varied across time points, which was to mimic the real case scenario as the change of the lab equipment and the sequencing technology due to the time change may change error rates.

In our simulation, we varied six variables, which are false positive rate, false negative rate, missing rate, $u$ controlling unobserved nodes, $a$ controlling # mutations, and varying error rates. We list all the varying numbers of these six variables and their defualt setting in **Supplemental Table S1**.

### 3 Determining the threshold for eliminating clusters

The FP and FN rates may change after we eliminate a subclone because the cells belonging to the subclone will be ascribed to other subclones in the same time point. A wrong elimination of the subclone will lead to an increase of the FP and FN rates. Thus to check whether our elimination is legitimate, we set up the max FP and FN rates for each time point as a threshold. We avoid making any constant threshold as they may vary from time point to time point. Instead, we calculate such a threshold from the data. Specifically, based on the simulated data set whose ground truth FP and FN rates for each time point is known, we observed that BnpC's inferred error rate for both FP and FN rates are about 0.8 of the ground truth error rate, seen in **Supplemental Fig. S18** and **Supplemental Fig. S19**. Therefore, we calculated the *threshold* of FP for each time point as *max* (inferred FP rates from all BnpC runs) / 0.8 for each time point. Similarly, we calculated the *threshold* of FN for each time point as *max* (inferred FN rates from all BnpC runs) / 0.8 for each time point. A subclone will not be eliminated if the elimination results in either FP or FN rate exceeding their corresponding threshold.

### 4 Illustration of Tree Inference Algorithm

In the main text, we described with an illustration how to infer such a tree when there is only one layer of unobserved nodes between time $k$ and $k+1$. Here we give an illustration of inferring a tree that has more than a layer of unobserved nodes in between two time points. Suppose on time $k$, scLongTree inferred two subclones: N1 ({1}) and N2 ({2, 3}). Here the numbers in the parenthesis are the mutations that each subclone's genotype had. On time $k+1$, scLongTree inferred five subclones: N3 ({1, 4, 5}), N4 ({1, 4, 6, 7}), N5 ({1, 4, 6}), N6 ({2, 3, 8, 9}) and N7 ({2, 3, 8}).

As shown in **Supplemental Fig. S4**, in the first round of the tree inference algorithm, we aimed at inferring the first layer of the unobserved nodes in between time $k$ and $k+1$. There were two inputs to the algorithm. The first input was a hash table in which the key was the mutations and the value was the subclones that contained the mutations. Here we skip the step of merging keys with the same value, and suppose the keys have already been merged. As it can be seen, N6 and N7 shared mutations {2, 3, 8} (first entry of the hash table); N3, N4 and N5 shared mutations {1, 4} (second entry), and N4 and N5 shared mutation {6} (third entry). All the rest of the entries had only one subclone in the value. We will illustrate the whole process in four steps, as shown in **Supplemental Fig. S4**. Step 1 eliminates those entries with only one subclone, as well as find the maximum set of mutation in a key for the corresponding value in an entry. Step 2 decreasingly sorts the entries by the number of subclones in the value. Step 3 checks if the top entry's key is a superset of any subclone's mutation set in time $k$. Step 4 will add the unobserved node, and connect it with the subclone in time $k$. Step 4 will also remove from the hash table all those subclones that appear in the values of the entry that is used to build the unobserved node. It will also remove all entries which have only one node in the value, and the entry that is used to build the unobserved node. In the illustration shown in **Supplemental Fig. S4**, **step 1** removed the entries that had only one subclone in the value. In addition, step 1 also replaced the remaining keys with the maximum subset of mutations contained by all subclones in the corresponding values. Thus the key of value N4 and N5 was replaced with mutations {1, 4, 6} since these are the mutations shared by N4 and N5. After **Step 2**, the entries were decreasingly sorted by the number of subclones in the value. **step 3** then checked whether the first entry's key, {1, 4}, was a superset of one of the subclones in the previous time point, $k$. **Step 4** then identified the unobserved nodes. Since set {1, 4} was a superset of {1} which was the mutation contained by N1 at time $k$, the first unobserved node was identified, called U1, and U1's mutation set was {1, 4}. The algorithm then connected N1 and U1, and the edge between the two had a new mutation 4. In this step (step 4), we also removed N3, N4 and N5 from the other entries' values because they will be the descendants of U1. Since the third entry had only N4 and N5 in the values, after the removal of the values, this entry became empty and thus was eliminated. We also eliminated the first entry with key {1, 4} since this entry had already been processed. We then repeated step 3 which searched for a key whose set of mutations was a superset of any of the subclones at time $k$, and step 4 which constructed the unobserved subclone. We found another unobserved subclone, called U2, whose mutations were {2, 3, 8}. We connected U2 with its parent node N2, and annotated a new mutation 8 on their edge. After this, the hash table became empty.

In the second round of the tree inference algorithm (**Supplemental Fig. S5**), we searched for the second layer of the unobserved nodes in addition to those found in the first round. The second layer of the unobserved nodes will be placed below the nodes in time $k$ or the first layer of the unobserved nodes. In

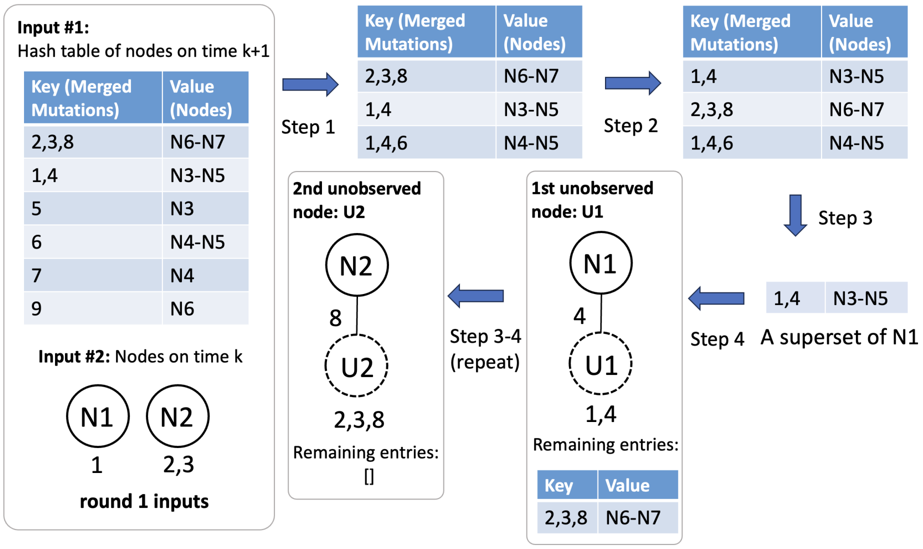


Supplemental Fig. S4: Illustration of the process of searching for the unobserved subclone in the first layer. The first input was a hash table whose keys were the merged mutations and values were the subclones at time $k+1$ containing the merged mutations. The second input was the subclones detected in time $k$. We put the mutations that these subclones contained under the circles. Outputs were the unobserved nodes in dashed circles (U1 and U2), as well as the annotations on the edges as the new mutations.

this round, the first input was exactly the same as that of the first round. The second input contained not only the subclones detected in time $k$, but also those unobserved subclones identified in the first round. Thus there were in total four nodes in the second input, which were N1, N2, U1 and U2. We first remove the entries whose keys were exactly the same as one of the nodes in input #2. After step 1 and 2, we had only one entry remaining in the hash table, whose key was $\{1,\ 4,\ 6\}$. In step 3, since the remaining entry's key $\{1,\ 4,\ 6\}$ was a superset of the mutations of both N1 and U1, we selected U1 as the priority was given to the unobserved node. In this way, we identified an unobserved node, called U3 whose parent node was U1, and on their edge was a new mutation 6. The algorithm stopped as this last entry was eliminated after it was processed, and the hash table became empty.

At last, for each subclone in time $k+1$, we connected it with the unobserved node whose value of the entry contained this subclone. We gave priority to the unobserved node in the second layer. For example, subclones N4 and N5 appeared in the values of both $\{1,\ 4\}$ which U1 has, and $\{1,\ 4,\ 6\}$ which U3 has. Since $\{1,\ 4,\ 6\}$ was processed and used to construct the unobserved node U3 in the second layer, whereas $\{1,\ 4\}$ was processed in the first layer, we connected

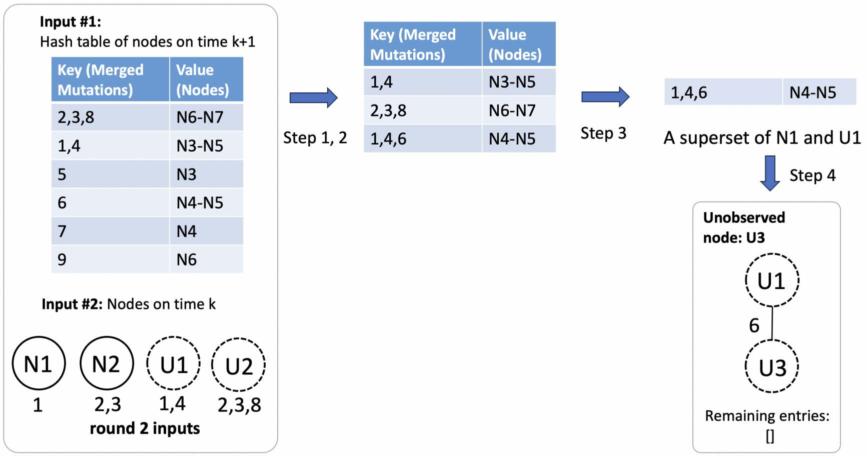


Supplemental Fig. S5: Illustration of the process of searching for the unobserved subclone in the second layer. The first input was a hash table whose keys were the merged mutations and values were the subclones at time $k + 1$ containing the merged mutations. The second input was the subclones detected in time $k$, as well as those unobserved nodes detected in the first layer. Here the second inputs were N1, N2, U1, and U2. Output was the new unobserved node U3 in dashed circle that had a new mutation 6 and was a child node of U1.

N4 and N5 with U3 instead of U1. The final longitudinal subclone tree for these two time points is shown in **Supplemental Fig. S6**.

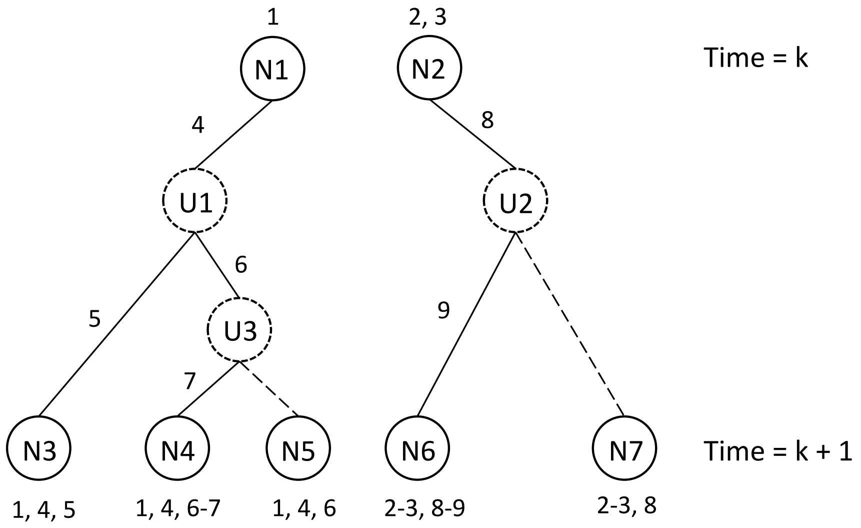


Supplemental Fig. S6: The inferred longitudinal subclonal tree between time point $k$ and $k + 1$. Unobserved nodes are in dashed circles. Edges without new or back mutations are in dashed lines. Mutations each subclone has are annotated on the top or bottom of the circles.

### 5 Reassigning cells to new clusters

When a cluster is eliminated, we reassign its cells to the other clusters in the same time point.

Specifically, for each cell $i$ in time point $k$, we use the following equation to calculate its likelihood of belonging to a subclone $l$ , $L_j(i|g)$, in terms of one mutation $j$. In this calculation, we do not consider the mutations with missing data or unknown consensus genotype because these entries are not informative of the subclone that the cell belongs to.

$$L_j(i|g) = \begin{cases} 1-\alpha, & \text{if } D_{i,j} = 0, g_j = 0 \\ \alpha, & \text{if } D_{i,j} = 1, g_j = 0 \\ \beta, & \text{if } D_{i,j} = 0, g_j = 1 \\ 1-\beta, & \text{if } D_{i,j} = 1, g_j = 1 \end{cases}$$

We then calculate the log likelihood $\sum_j \log L_j(i|g)$ for all eligible mutations which do not have missing entry on cell $i$. We compute the average of the log likelihood by dividing $\sum_j \log L_j(i|g)$ by the number of eligible mutations for each cluster $l$, and select the cluster that has the highest likelihood to be the cluster that cell $i$ belongs to.

### 6 Running scLongTree, LACE, SiCloneFit, and SCITE for evaluation

#### 6.1 scLongTree:

We ran BnpC with default settings for getting the clustering results. BnpC's estimated FP and FN rates are used as the initial FP and FN rates in inferring the longitudinal tree. We set $k = 0$ to eliminate the back mutations. $k$ can be tuned according to the number $k$ set in the $k$-Dollo model.

#### 6.2 LACE:

LACE was executed with their default settings by passing only the D matrix as an input.

#### 6.3 SiCloneFit:

We used the following parameter/arguments for running SiCloneFit on all simulated datasets:

- -iter 100
- -fp 0.01
- -fn 0.2

- -r 10
- -df 0

##### 6.4 SCITE:

We used the following parameter/arguments for running SCITE on all simulated datasets:

- -r 1
- -l 1565000
- -fd 0.01
- -ad 0.2
- -max_treelist_size 1

##### 6.5 SCG:

SCG was run using a three-state error model with the following hyperparameters. The gamma priors for the false-positive, false-negative, and mutation states were set to (9.99, 0.01, $10^{-15}$), (2.5, 7.5, $10^{-15}$), and ($10^{-15}$, $10^{-15}$, 1), respectively. The state priors were set to [1, 1, $10^{-15}$], and the Dirichlet process concentration parameter was set to 1 for all simulated datasets.

##### 6.6 RobustClone:

RobustClone was executed with the default pipeline:MATLAB RPCA on the $D$ matrix (0/1/3) followed by the provided R clustering/MST scripts—using only the $D$ matrix as input.

RobustClone outputs an undirected minimum-spanning tree of subclones without explicit ancestral/parallel semantics, so those labels cannot be assigned without extra assumptions. We therefore restrict RobustClone to same-edge pairs obtained from its clone–mutation mapping and compute accuracy on the intersection of mutation IDs present in both outputs. Consequently, the RobustClone bars in **Fig. 2** reflect same-edge accuracy only.

#### 7 Selecting the best BnpC run

For each dataset and replicate, BnpC was executed five times, producing alternative clustering solutions. To determine which run best explains the observed single-cell mutation matrix, we evaluated each run using a likelihood model consistent with the false-positive (FP) and false-negative (FN) error rates inferred by BnpC.

For a given run, let $D_{i,j} \in \{0, 1\}$ denote the observed mutation state of cell $i$ at site $j$, and let $g_{l,j} \in \{0, 1\}$ be the inferred genotype of cluster $l$ at the same site. Missing entries in the observed data were excluded. For each non-missing entry, the probability of observing $D_{i,j}$ given the cluster genotype was

$$P(D_{i,j} \mid g_{l,j}) = \begin{cases} 1-\alpha, & D_{i,j}=0,\ g_{l,j}=0, \\ \alpha, & D_{i,j}=1,\ g_{l,j}=0, \\ \beta, & D_{i,j}=0,\ g_{l,j}=1, \\ 1-\beta, & D_{i,j}=1,\ g_{l,j}=1, \end{cases}$$

where $\alpha$ and $\beta$ are the inferred FP and FN rates by BnpC, respectively.

For each $(i,j)$ pair, we computed the log-likelihood contribution $\log P(D_{i,j} \mid g_{l,j})$. The total log-likelihood for a run is the sum of these contributions across all cells and mutations.

Because BnpC outputs results separately for each time point, we computed the overall log-likelihood for each run by summing the log-likelihoods from all time points within the same run. Among the five BnpC runs, the run with the highest total log-likelihood was selected as the best solution for that dataset and run.

### 8 Evaluation metrics

#### 8.1 Pairwise SNV accuracy:

We defined three different scenarios to calculate the pairwise SNV accuracy:

- Parallel SNVs $ps$: A pair of SNVs whose edges are in parallel with each other.
- Ancestral SNVs $as$: A pair of SNVs whose edges are in an ancestral-descendant relationship.
- SNVs on the same edge $is$: A pair of SNVs on the same edge.

Given each of the three inferred pairwise SNVs set $(I_{ps}, I_{as}, I_{is})$, we compare them with the true pairwise SNVs set $(GT_{ps}, GT_{as}, GT_{is})$, and compute their intersection sets' sizes. In this way, we count the number of correctly inferred pairwise SNVs. We calculate the pairwise SNV accuracy using the following equation.

$$Accuracy = \frac{|I_{ps} \cap GT_{ps}| + |I_{as} \cap GT_{as}| + |I_{is} \cap GT_{is}|}{|GT_{ps} \cup GT_{as} \cup GT_{is}|}$$

#### 8.2 SNV placement accuracy:

We measured the accuracy of the placement of SNVs in each time point by the time point the SNV is placed. On a certain time point $k$, we define $TP_k$ as the number of SNVs that are placed on $k$ while the ground truth also places it on

$k$. We define $FP_k$ as the number of SNVs that are placed on $k$ while the ground truth does not place it on $k$. We define $FN_k$ as the number of SNVs that are not placed on $k$ while the ground truth places it on $k$. We then define precision and recall at a certain time point $k$ as follows.

- Precision $p_k = \frac{TP_k}{TP_k+FP_k}$
- Recall $r_k = \frac{TP_k}{TP_k+FN_k}$

We then average the precision and recall from all time points, the result of which are the final precision $p$ and final recall $r$. Finally, we calculate the accuracy as the harmonic mean of $p$ and $r$, i.e., $\frac{2pr}{p+r}$.

### 9 Determining SCITE and SiCloneFit's SNV placement accuracy

#### 9.1 SCITE's SNV placement accuracy

SCITE does not use time points during inference, so we assign time points post hoc using ground truth. We first construct a cell→timepoint map from the ground-truth data. For each SCITE mutation node, we identify its associated cells and assign the node a time point by majority vote over the *ground-truth* time points of those cells. Because SCITE nodes are not annotated by SNV, each ground-truth SNV is linked to a SCITE node by maximizing the Jaccard overlap between the SNV's true carrier-cell set and each node's cell set; only matches whose best similarity exceeds a preset threshold are considered for evaluation. The SNV's *predicted* time point is the majority-vote time point of its matched node, while its *ground truth* time point is the majority vote over the ground-truth time points of its true carrier cells. We report *accuracy* as the fraction of SNVs with confident matches whose predicted time point equals ground truth.

#### 9.2 SiCloneFit's SNV placement accuracy

To determine SNV placement for SiCloneFit, we use a cluster-based strategy that is robust to single-cell noise. SiCloneFit infers clusters of cells and their genotypes, while the true time point of each cell is known from the ground truth. For each inferred cluster, we examine the ground-truth time points of its member cells and assign the cluster to the time point that contains the majority of its cells (ties are broken by choosing the earliest time point). In this way, each inferred cluster is mapped to a single time point based on where most of its cells originate in the ground truth.

After assigning clusters to time points, we determine which mutations belong to each cluster using the following rule : A mutation is considered present in a cluster if more than half of the cells in that cluster carry the mutation according to the inferred genotypes. All mutations present in clusters assigned to the same time point are then pooled together. We process time points in chronological order and assign each mutation to the earliest time point at which it newly appears compared to previous time points. This inferred mutation time point is then compared with the ground-truth mutation time point to evaluate SNV placement accuracy.

### 10 Supplemental Tables

| | |
|---|---|
| **False positive rate (FP)** | 0.001, 0.01(d), 0.05 |
| **False negative rate (FN)** | 0.1, 0.2(d), 0.3, 0.4 |
| **Missing rate (MR)** | 0.2(d), 0.3 |
| $u$ **controlling unobserved nodes** | 0, 0.1, 0.2(d), 0.3 |
| $a$ **controlling # mutations** | 1.7(d) (50), 3.4 (85), 5.1 (140) |
| **Varying error rates across time points:**<br>**1. constant**<br>**2. less varied**<br>**3. more varied** | 1. FP = 0.01, FN = 0.2, MR = 0.2 (d)<br>2. FP = [0.01, 0.03], FN = [0.2, 0.3], MR = 0.2<br>3. FP = [0.001 , 0.01, 0.03, 0.05], FN = [0.1, 0.2, 0.3, 0.4], MR = [0.2, 0.3] |

Supplemental Table S1: A list of the variables varied in the simulated datasets. Each line has a varying variable (first column) with the values in the second column. The default value is denoted by “(d)” on its right.

| **Mutation constant** $a$ | **1.7 (50)** | **3.4 (85)** | **5.1 (140)** |
|---|---|---|---|
| **scLongTree including BnpC’s runtime (s)** | 27433 | 21482 | 42095 |
| **scLongTree not including BnpC’s runtime (s)** | 6 | 6 | 7 |
| **LACE (s)** | 7802 | 8492 | > 24 hrs |
| **SCITE (s)** | 525 | 325 | 914 |
| **SiCloneFit (s)** | 13155 | 16835 | 39088 |

Supplemental Table S2: Median of running time in seconds for varying variable $a$ for scLongTree including BnpC’s runtime, scLongTree not including BnpC’s runtime, LACE, SCITE and SiCloneFit.

### 11 Supplemental Figures

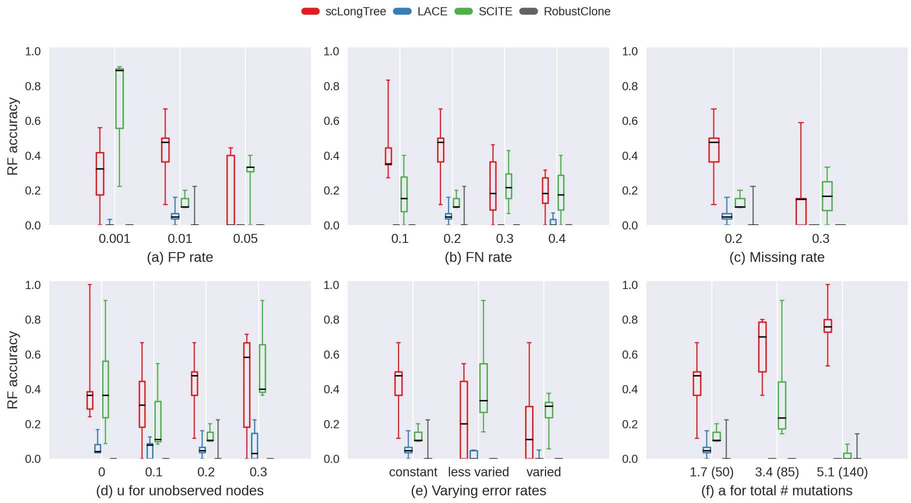


Supplemental Fig. S7: Comparison of Robinson-Foulds (RF) accuracy between the inferred tree structure and the ground truth tree among scLongTree (red), LACE (blue), SCITE (green) and RobustClone (grey). SiCloneFit's RF are most of the time 0 due to the lack of clusters and thus are not shown in this figure.

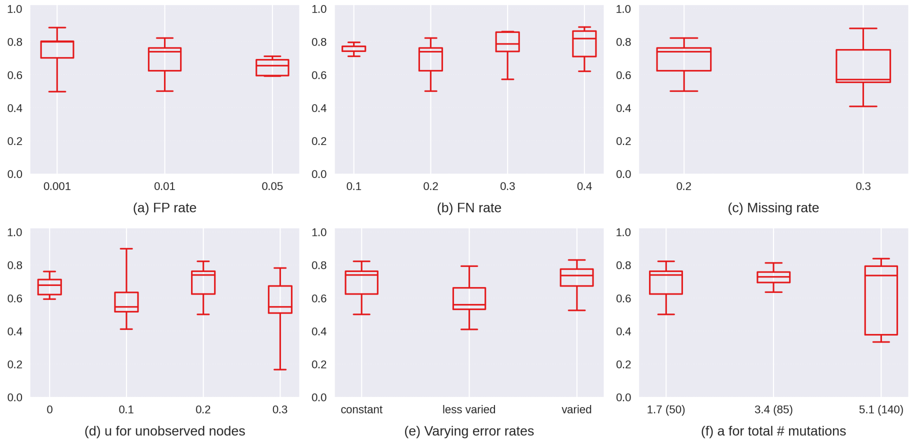


Supplemental Fig. S8: ScLongTree's imputation accuracy across all variables.

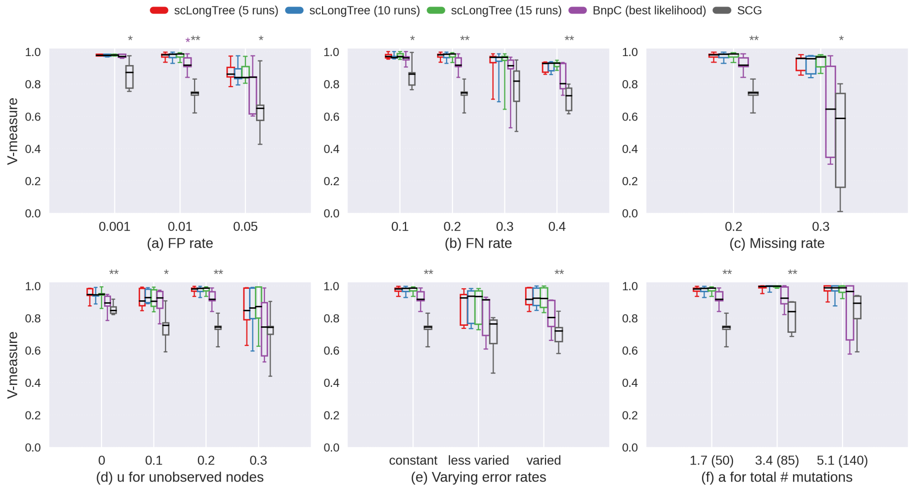


Supplemental Fig. S9: Comparison of cell clustering accuracy (V-measure) for scLongTree with 5 (red), 10 (blue) and 15 (green) runs of BnpC, the BnpC alone with the best likelihood (purple), and SCG (black). Statistical significance of the Student t-test is indicated by asterisks: * ($p < 0.01$), and ** ($p < 0.05$).

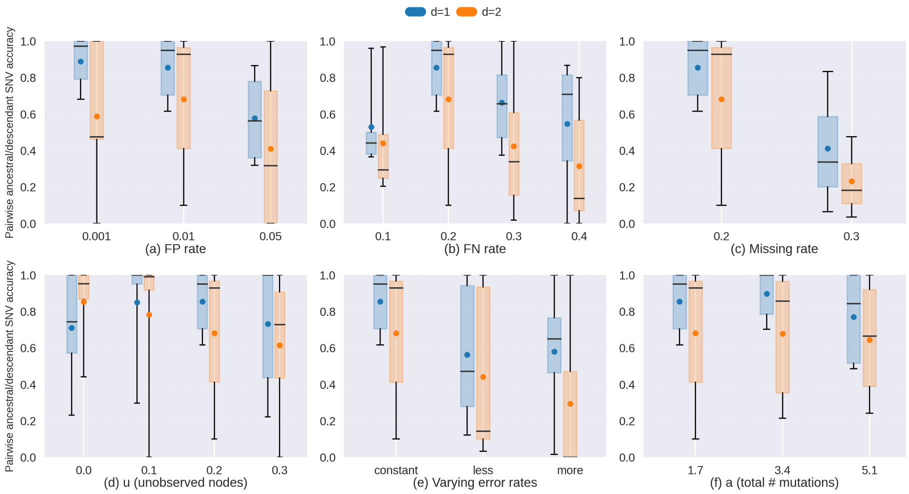


Supplemental Fig. S10: ScLongTree's pairwise SNV accuracy in terms of the parent-child (d=1, blue bars) and grandparent-grandchild (d=2, red bars) relationship for the simulated dataset.

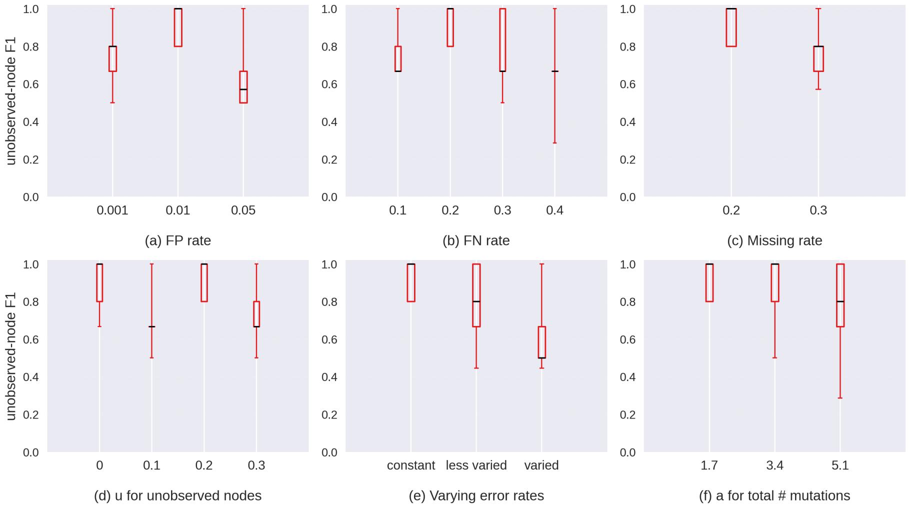


Supplemental Fig. S11: ScLongTree's F1 score of the inferred unobserved nodes for the simulated dataset. A match of an inferred unobserved node and a ground truth unobserved node requires that their observed parent or daughter nodes share a Jaccard Index $\geq 0.75$ based on the cells they contain, or that they both are the daughter nodes of the root.

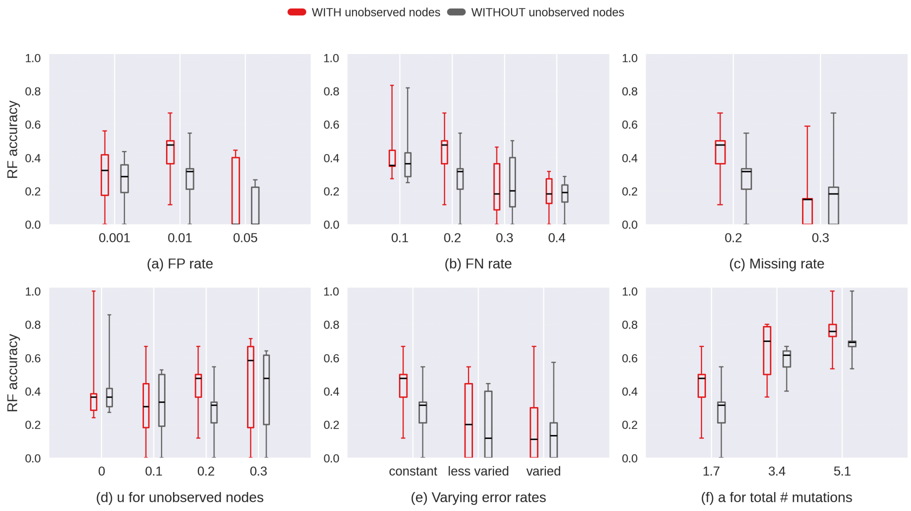


Supplemental Fig. S12: ScLongTree's Robinson Foulds score with (in red bars) and without (in black bars) the unobserved nodes for the simulated dataset.

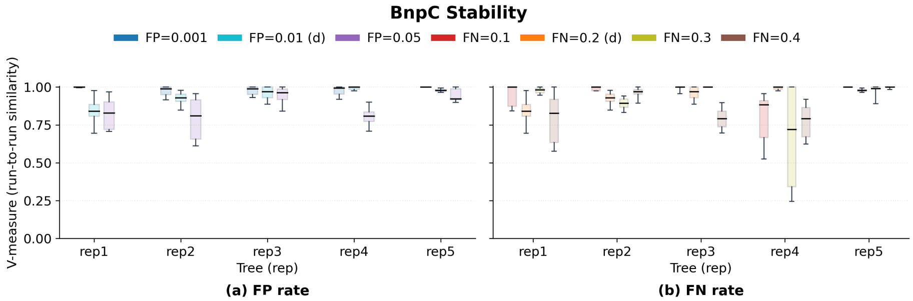


Supplemental Fig. S13: Run-to-run similarity of BnpC to evaluate BnpC's stability for a) varying FP rate and b) varying FN rate.

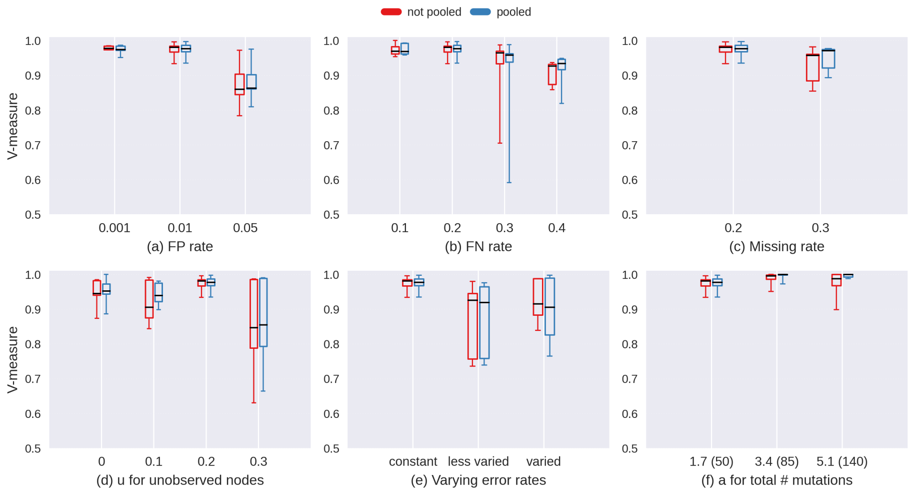


Supplemental Fig. S14: Comparison of the V-measure of scLongTree pooling (blue) and not pooling (red) cells from all time points.

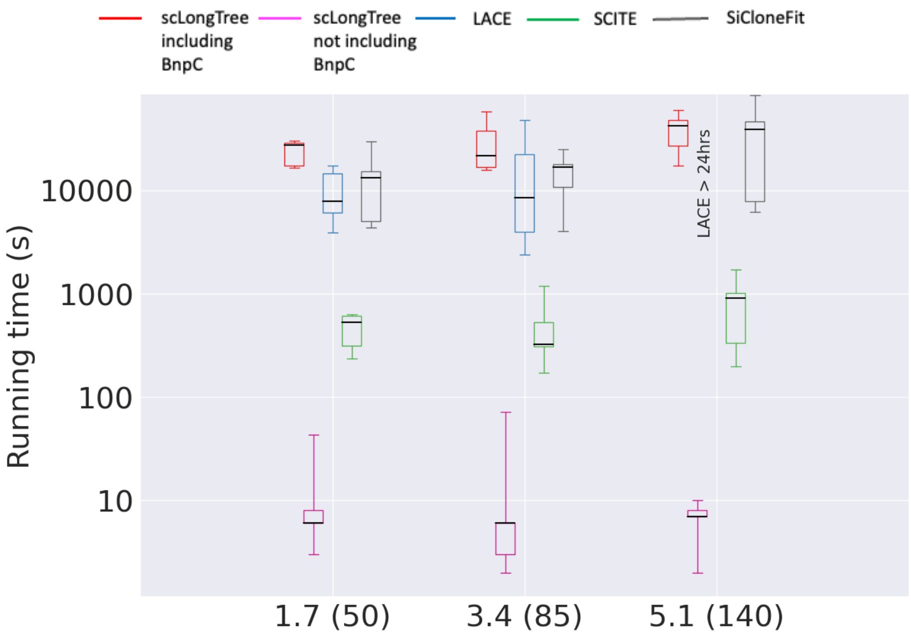


Supplemental Fig. S15: Boxplots are shown for running time in seconds for varying variable $a$ for scLongTree including BnpC's running time (red), scLongTree not including BnpC's running time (pink), LACE (blue), SCITE (green) and SiCloneFit (black). Median of each boxplot was highlighted in a black horizontal line.

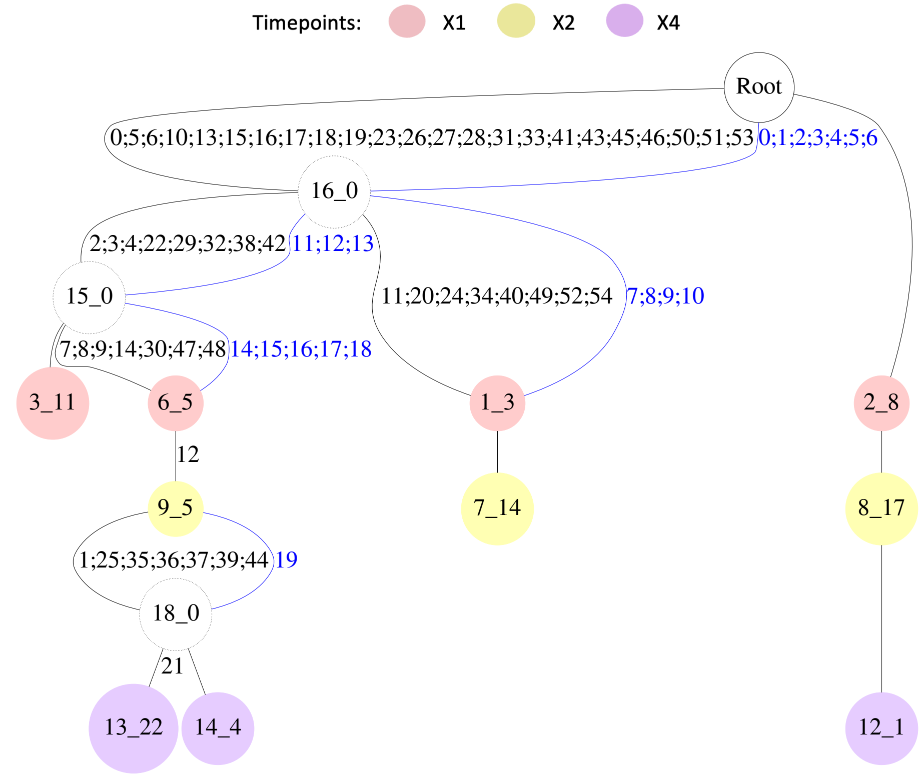


Supplemental Fig. S16: ScLongTree's inferred longitudinal subclonal tree for SA501 given the large SA501 dataset (55 mutations). The black numbers on the black edges represent the 55 mutations. The corresponding small SA501 dataset (20 mutations) in its own index system is shown in blue numbers on blue edges.

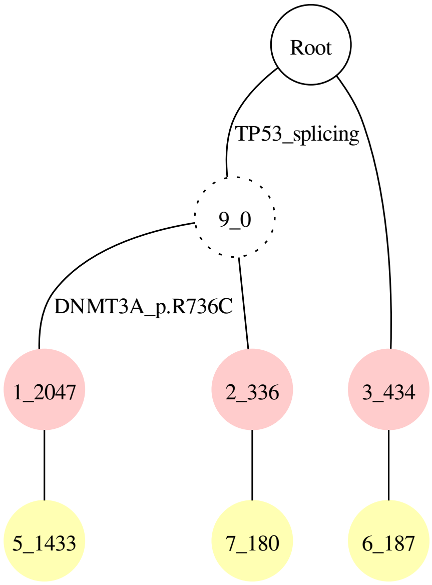


Supplemental Fig. S17: ScLongTree inferred longitudinal subclonal tree for real dataset AML107.

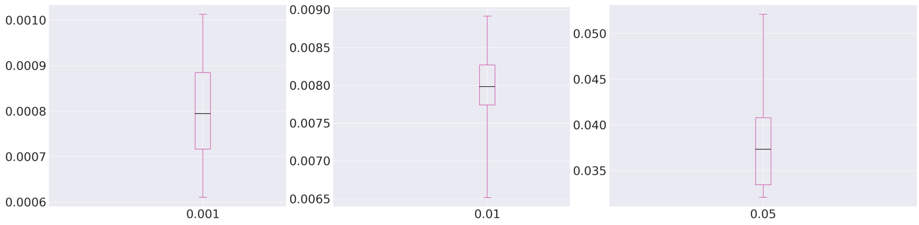


(a) FP = 0.001 (b) FP = 0.01 (c) FP = 0.05

Supplemental Fig. S18: Boxplots are shown for various simulated FP rates where Y-axis has the BnpC estimated FP rates and X-axis has the simulated FP rates. The BnpC estimated FP rates are approximately 0.8 of the true values.

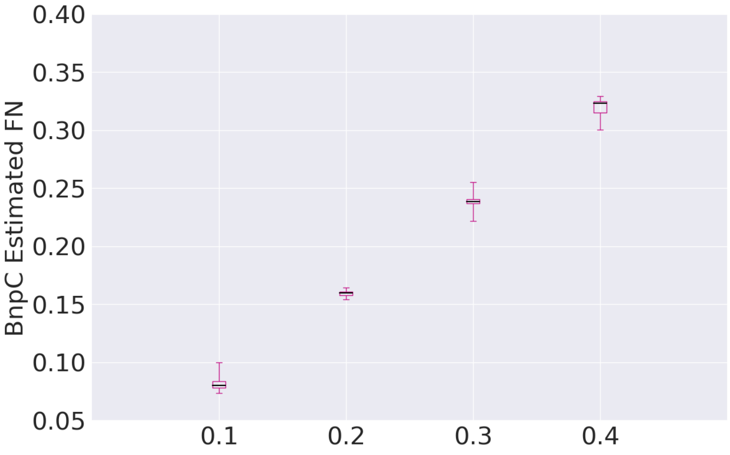


Supplemental Fig. S19: Boxplots are shown for various simulated FN rates where Y-axis has the BnpC estimated FN rates and X-axis has the simulated FN rates. The BnpC estimated FN rates are approximately 0.8 of the true values.

### 12 Supplemental Algorithm S1

**Algorithm 1** scLongTree Pseudocode

**Inputs:**
$D$ = input noisy genotype matrix
$B_t$ = BnpC clusters for each time point from all runs
$G$ = BnpC cluster genotype matrix
$n$ = No. of BnpC runs
$\alpha_t, \beta_t$ = BnpC estimated FP and FN rates for each time point $t$
**Output:**
$Tree$ = Longitudinal Tree with mutations on its edges and cells in each clone
**Parameters:**
$\hat{\alpha_t}$ = FP threshold for each time point
$\hat{\beta_t}$ = FN threshold for each time point
$\gamma_t$ = Missing rate for each time point
$\alpha_i, \beta_i, Tree_i, TP_i$ = Final FP, FN rates, Tree and Tree probability from each BnpC run $i$.

**for** $i \epsilon n$ **do**
  $P_{B_i}$ = Calculate cluster probabilities using $D, G, \alpha_t i, \beta_t i, \gamma_t$ and Eq. 1. Probabilities are then sorted in ascending order.
  tree_prob = $\prod$P$_{B_i}$
  $Tree$ = Infer Tree from the clustering results.
  **for** $c \, \epsilon \, P_{B_i}$ **do**
    Re-calculate $\alpha_t i$ and $\beta_t i$ after dropping subclone $c$ and reassigining its cells to another cluster with *MaxLikelihood*.
    **if** $\alpha_t i > \hat{\alpha_t} \vee \beta_t i > \hat{\beta_t}$ **then** don't eliminate $c$, retain its cells.
    **end if**
    $Tree$ = Infer new $Tree$.
    Re-calculate new_tree_prob with updated $P_{B_i}$ for clusters with reassigned cells.
    **if** new_tree_prob > tree_prob **then** tree_prob = new_tree_prob. Save the $Tree$. Update $P_{B_i}$ with new cluster probabilities and **continue**.
    **end if**
  **end for**
  $TP_i$ = tree_prob of the $Tree$ with the highest probability from this run.
  $Tree_i$ = Tree after correcting parallel and back mutations using $k$-Dollo model.
  $\alpha_i, \beta_i$ = Final FP, FN rates after parallel and back mutation correction.
**end for**
Eliminate the BnpC runs having $\alpha_i$ or $\beta_i$ > its (median + standard deviation) from all runs. From the remaining BnpC runs we selected the $Tree$ with max $(TP_i)$.
